# Genetic polymorphisms of Leukocyte Immunoglobulin-Like Receptor B3 (*LILRB3*) gene in African American kidney transplant recipients are associated with post-transplant graft failure

**DOI:** 10.1101/2024.02.21.581383

**Authors:** Zeguo Sun, Zhengzi Yi, Chengguo Wei, Wenlin Wang, Paolo Cravedi, Fasika Tedla, Stephen C. Ward, Evren Azeloglu, Daniel R. Schrider, Yun Li, Sumaria Ali, Tianyuan Ren, Shun Liu, Deguang Liang, Jia Fu, Tong Liu, Hong Li, Caixia Xi, Thi Ha Vy, Gohar Mosoyan, Quan Sun, Ashwani Kumar, Zhongyang Zhang, Samira Farouk, Kirk Campell, Jordi Ochando, Kyung Lee, Steve Coca, Jenny Xiang, Patti Connolly, Lorenzo Gallon, Robert Colvin, Madhav Menon, Girish Nadkarni, John C. He, Monica Kraft, Xuejun Jiang, Xuewu Zhang, Weiguo Zhang, Shu-hsia Chen, Peter Heeger, Weijia Zhang

## Abstract

**Background:** African American (AA) kidney transplant recipients exhibit a higher rate of graft loss compared to other racial and ethnic populations, highlighting the need to identify causative factors underlying this disparity.

**Method:** We analyzed RNA sequences of pretransplant whole blood from subjects followed in three kidney transplant cohorts to identify single nucleotide polymorphisms (SNPs) associated with death censored graft loss (DCGL). We employed a meta-analysis to uncover key transcriptional signatures and pathways associated with the identified SNPs and used single cell RNA to define cellular specificity. We characterized SNP functions using *in vitro* immunological and survival assays and tested for associations between the identified SNPs and other immune-related diseases using a ∼30,100 subject, electronic health record (EHR)-linked database.

**Results:** We uncovered a cluster of four consecutive missense SNPs in the Leukocyte Immunoglobulin-Like Receptor B3 (*LILRB3*, a negative immune response regulator) gene that strongly associated with DCGL. This *LILRB3*-4SNPs cluster encodes missense mutations at amino acids 617-618 proximal to a SHP-1/2 phosphatase-binding ITIM motif. *LILRB3*-4SNPs is specifically enriched within subjects of AA ancestry (8.6% prevalence vs 2.3% in Hispanic and 0.1% in European populations), is not linked to *APOL1* G1/G2 alleles, and exhibited a strong association with DCGL. Analysis of PBMC and transplant biopsies from recipients with *LILRB3*-4SNPs showed evidence of enhanced adaptive immune responsiveness and ferroptosis-associated death in monocytes. Overexpression of the variant allele in THP-1 cells (macrophage line) induced augmented inflammation and ferroptosis, which were attenuated by a ferroptosis inhibitor, verifying a causal link. The *LILRB3*-4SNPs also associated with multiple systemic and organ-specific immune-related diseases in AAs, consistent with conferring a broadly relevant immune function.

**Conclusion:** the *LILRB3*-4SNPs represent a functionally important, distinct genetic risk factor for kidney transplant outcome and development/severity of other immune-related diseases in patients of AA ancestry. Pharmacological targeting of ferroptosis should be tested to prevent or treat these disease processes in AA recipients carrying *LILRB3*-4SNPs.

## Introduction

African American (AA) kidney transplant recipients face a higher risk of graft failure compared to individuals of European or other ancestries^1–3^. This elevated risk is has been validated in subjects with genetic markers of African ancestry ^4,5^ but is not attributable to higher/different HLA disparities compared to other groups ^1^. While several genome-wide association studies (GWAS) have identified genetic polymorphisms in HLA and non-HLA regions associated with post-transplant graft failure in a mixed population ^5,6^, whether non-HLA SNPs contribute to the reduced graft survival in these patients remains inadequately tested^3^.

While SNP arrays have been commonly used for genotyping in GWAS, these arrays do not adequately probe SNPs in exonic regions or in the under-represented populations and do not provide quantitative association analysis of variant alleles with diseases, all of which limit interpretation of findings based on this methodology. RNA sequencing (RNAseq) overcomes these deficiencies by enabling the simultaneous detection of coding variants and quantification of overall gene expression and allele-specific expression levels of genes. This capability makes RNAseq a complementary tool for identifying functional SNPs and related gene signatures associated with human diseases in a quantitative manner.

Herein, we employed RNA sequencing of pretransplant blood samples from kidney transplant recipients to investigate the functional genetic polymorphisms involved in post-transplant graft loss. Through this analysis, we identified a cluster of four consecutive missense single-nucleotide polymorphisms (SNPs) in the close proximity to the ITIM motif of the Leukocyte Immunoglobulin-Like Receptors family B3 gene (*LILRB3*, an inhibitory immune response regulatory gene), called *LILRB3*-4SNPs that strongly associated with death censored graft loss and other immune mediated disease processes specifically in patients of African ancestry. Through mechanistic studies linking the genotype to function we determined that *LILRB3*-4SNPs augment inflammation and ferroptosis, thereby identifying an unanticipated therapeutic target that could potentially improve clinical outcomes in these patients.

## Results

### Study Cohorts

We analyzed subjects from three kidney transplant cohorts, GoCAR^7,8^, CTOT19^9^, VericiDx, and one EHR-linked biobank cohort (BioMe)^10^ (**Figure 1** and **Supplementary Methods)** with demographic characteristics summarized in **Table S1.**

**Figure 1.**
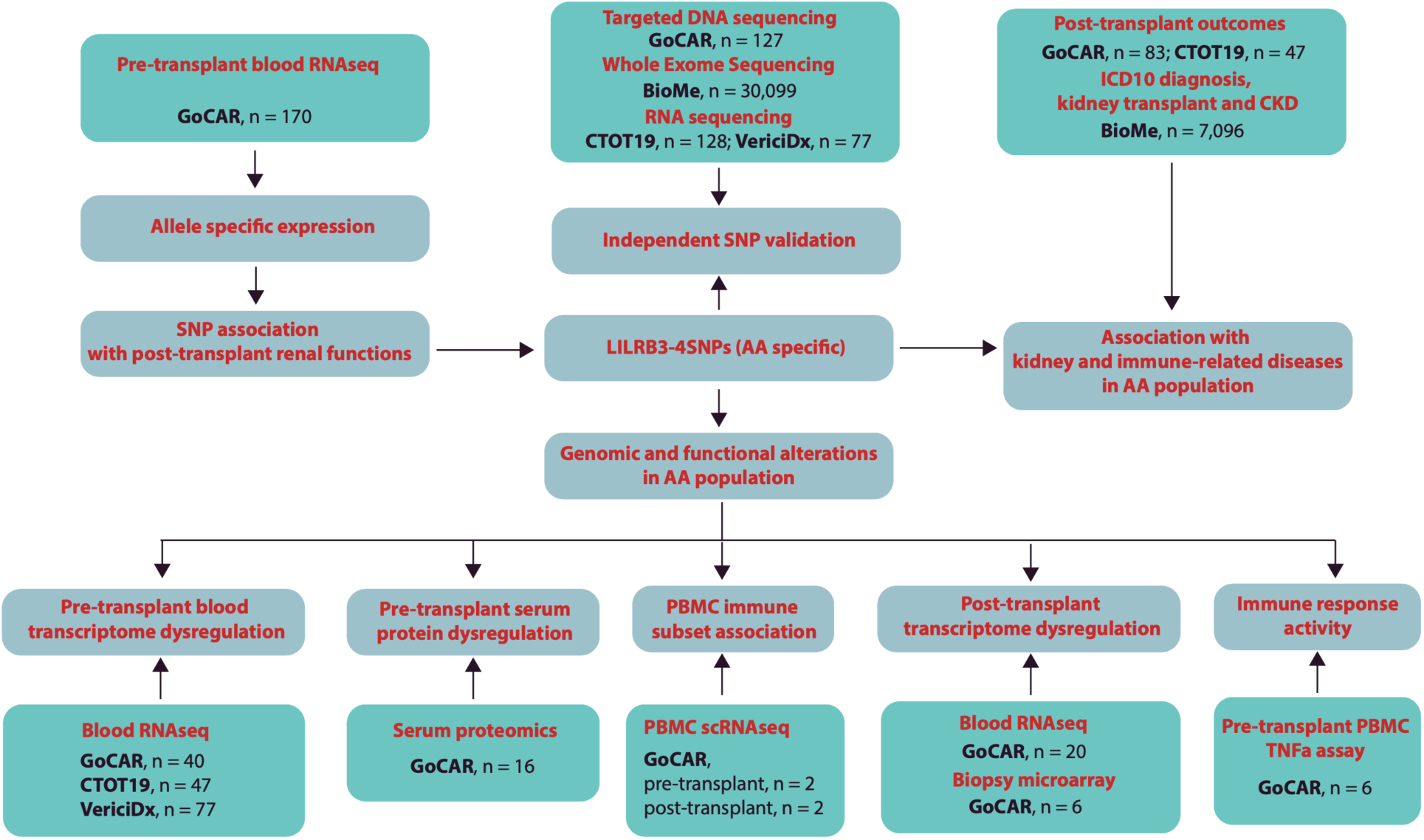
Cohort and study design. 3 independent kidney transplantation cohorts (GoCAR (n = 264, 83 AAs), CTOT19 (n=128, 47 AAs) and VericiDx (n = 77, 77 AAs)), and a large EHR linked BioMe biobank (n = 30,099, 7,096 AAs) were used in this study. Pretransplant blood RNAseq data was initially used to identify the AA-specific expressional SNPs (*LILRB3*-4SNPs) associated with post-transplant renal failure and validated by a targeted sequencing in GoCAR, RNAseq in CTOT19 and VericiDx and WES in BioMe. The genomic and functional alterations associated with *LILRB3*-4SNPs were investigated by multi-omics approaches (meta-analysis of RNA sequencing datasets of pretransplant blood samples from GoCAR, CTOT19 and VericiDx, mass spectrometry proteomics of pretransplant serum, single cell RNA sequencing of pre- and post-transplant PBMC, bulk RNAseq of post-transplant blood and biopsies and immune response assays on pretransplant PBMC in GoCAR cohort). The association of *LILRB3*-4SNPs with post-transplant kidney function and pathological lesions was assessed in GoCAR, CTOT19 and BioMe cohorts. Last, the association of *LILRB3*-4SNPs with the kidney diseases or other immune related diseases was evaluated in the large EHR-linked BioMe cohort.

We analyzed pre- or post-transplant specimens from the 264 subjects in GoCAR cohort and correlated results with clinical outcomes to identify, validate and functionally characterize the renal failure-associated SNPs (**Table S1** and **Table S2**). These 264 patients (71.6 % male) were composed of genetically-determined 83(31.4%) AAs, 82 (31.1%) Europeans and 86 (32.6%) Hispanics with an average age of 50.84 at the time of transplantation. The within 5-year graft loss rate is 19.7% in the entire cohort and 34.9% in AA population.

In the 128 recipient CTOT19 cohort with genetically-determined 47 (36.7%) AAs, 27 (21.1%) Europeans, 41 (32.0%) Hispanics, and 13 individuals from other ethnic backgrounds. We used RNAseq to validate the expression of SNPs identified from GoCAR cohort and validate correlations with 2-year transplant outcomes, and combined with samples from the other two transplant cohorts assessed SNP-associated gene expression dysregulation.

In the VericiDx cohort, 77 genetically-determined African American recipients with RNAseq data were combined with other two cohorts to identify SNP associated gene expression dysregulation. The relatively short follow-up duration for many patients in this cohort did not permit validation of outcome associations.

In the EHR-linked biobank cohort (BioMe), 30,099 patients with WES data and self-reported race information were used to validate the identified SNPs and estimate the prevalence or the minor allele frequency (MAF). 7,096 African American patients were used to investigate the association of the identified AA-specific SNPs with the severity of kidney diseases/transplant and other diseases defined by ICD-10 code.

### Genome wide screening of allele specific expression of transplant recipients in GoCAR

We developed a bioinformatic pipeline to quantify the expression of exonic SNPs within the GoCAR cohort (**Figure S1** and **Methods**). From genotyping arrays (n=588)^11^ and pretransplant RNAseq (n=170) ^8^ data in GoCAR cohort, 410,855 exonic SNPs were extracted for expression analysis on RNAseq data (n=170) ^8^ using the ASEReadCounter tool of the GATK toolkit^12^. At each SNP, we quantified the alternative allele expression fraction (AEF) as the fraction of the reads supporting alternative allele among all the reads supporting both alternative allele and reference allele of the SNP (see **Methods**). Within these exonic SNPs, 1,438 SNPs exhibited significant associations of AEF with death-censored graft loss (DCGL) with a threshold of nominal P ≤ 0.05 (**Figure 2A** and **Table S3**). This list was subsequently refined to 295 after being adjusted by donor status, HLA mismatch, induction therapy, and genetic race ^11^. These refined SNPs were situated within 255 distinct genes, displaying a significant enrichment in immune-related functionalities, including neutrophil degranulation, macrophage activation, and interferon gamma response (**Figure 2B** and **Table S4**), Upon tallying the count of significant SNPs within each gene region, multiple members of the Leukocyte Immunoglobulin-Like Receptor (LILR) gene family (LILRA2, LILRB1, *LILRB3* and LILRA1 with immune regulatory function^13^) occupied prominent positions within the top 20 genes (**Figure 2C**).

**Figure 2.**
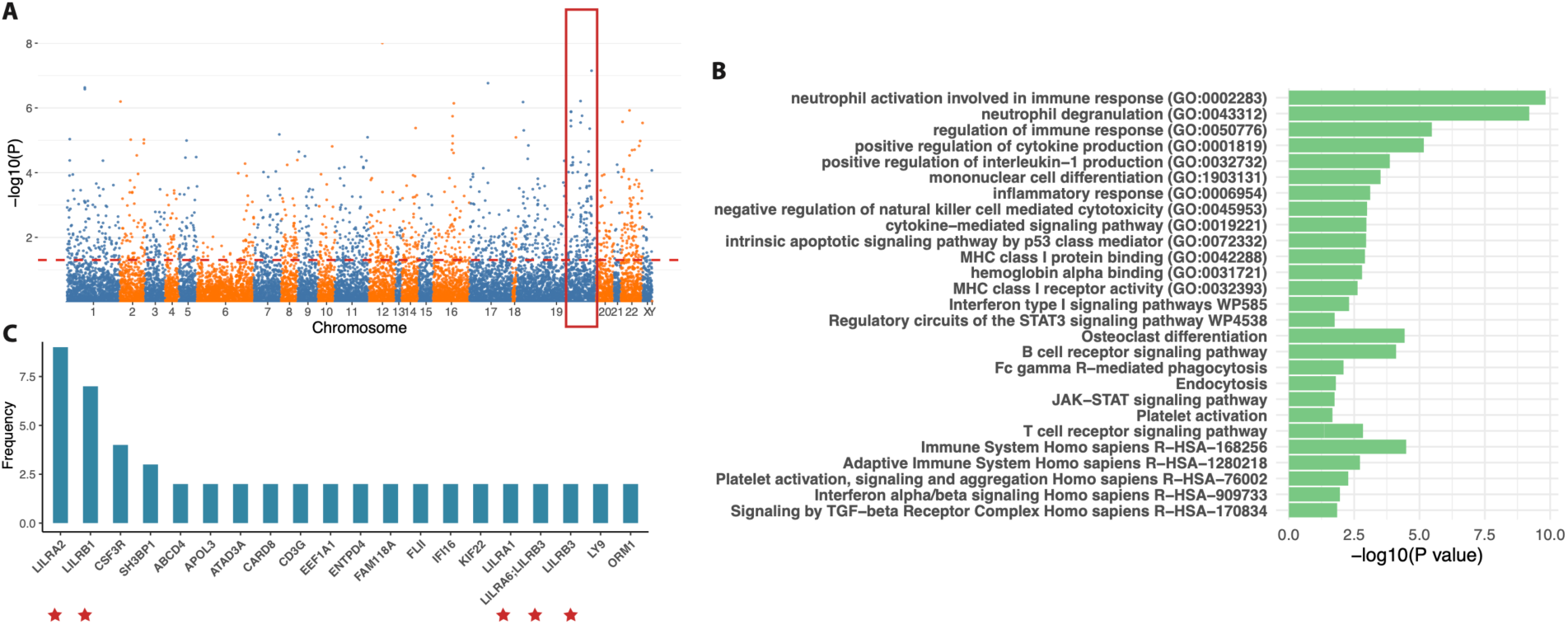
The association of expressional SNPs (eSNPs) with graft loss in GoCAR cohort. **A**) Manhattan plot of the eSNPs associated with DCGL; **B**) The enriched functions of the genes harboring significant eSNPs (P <= 0.05); **C**) The occurrence of significant eSNPs in the top 20 recurrent genes.

### Identification of a cluster of 4 consecutive missense SNPs in *LILRB3* (*LILRB3*-4SNPs) as a distinct, strong risk factor for post-transplant renal failure in GoCAR AA recipients

Due to the pivotal role of LILR genes as regulators of immune responses and their established connection with autoimmune diseases^13^, our focus gravitated towards discerning significant SNPs within the LILR gene families. Of notable interest was SNP (rs549267286 located at chr19:54721007 of hg19) – a missense SNP with high prevalence in the AA ancestry exhibited an intriguing correlation with DCGL (**Figure 3A**). However, upon an inspection of the read alignments covering rs549267286, we found this SNP is situated within a cluster of four highly-linked consecutive missense SNPs positioned at chr19:54721006-54721009 (rs139094141, rs549267286, rs567676351, and rs761515451, a region that we named *LILRB3*-4SNPs) (**Figure3 B** and **C** and **Figure S2**). Two adjacent synonymous SNPs (rs113314747 and rs60566950) within the *LILRB3* exon were also found to be in high LD with those 4 consecutive SNPs (**Figure3 B** and **C** and **Figure S2**). The remaining three SNPs within this cluster were not imputed via SNP array data or identified through RNA-seq genotyping, potentially attributed to the computational challenges for consecutive SNPs calling ^12^. *LILRB3*-4SNPs results in amino acid substitution specifically, 617-618 Glu-Pro (EP)-> Pro-Ser (PS). This substitution occurs in the linker region between ITIM3 and ITIM4 of *LILRB3*, raising the possibility that it could impact the binding strength of *LILRB3* to the SHP-1/2 protein^14^. A model of the complex between SHP2 and the ITIM3-linker-ITIM4 by AlphaFold suggest that the EP->PS mutation may alter the interactions between the ITIMs with SH2 domain of SHP2 ^15,16^ (**Figure 3D**). Worth noting is the conservation of the amino acid 617E across various *LILRB3* orthologs from diverse organisms (**Figure 3E**).

**Figure 3.**
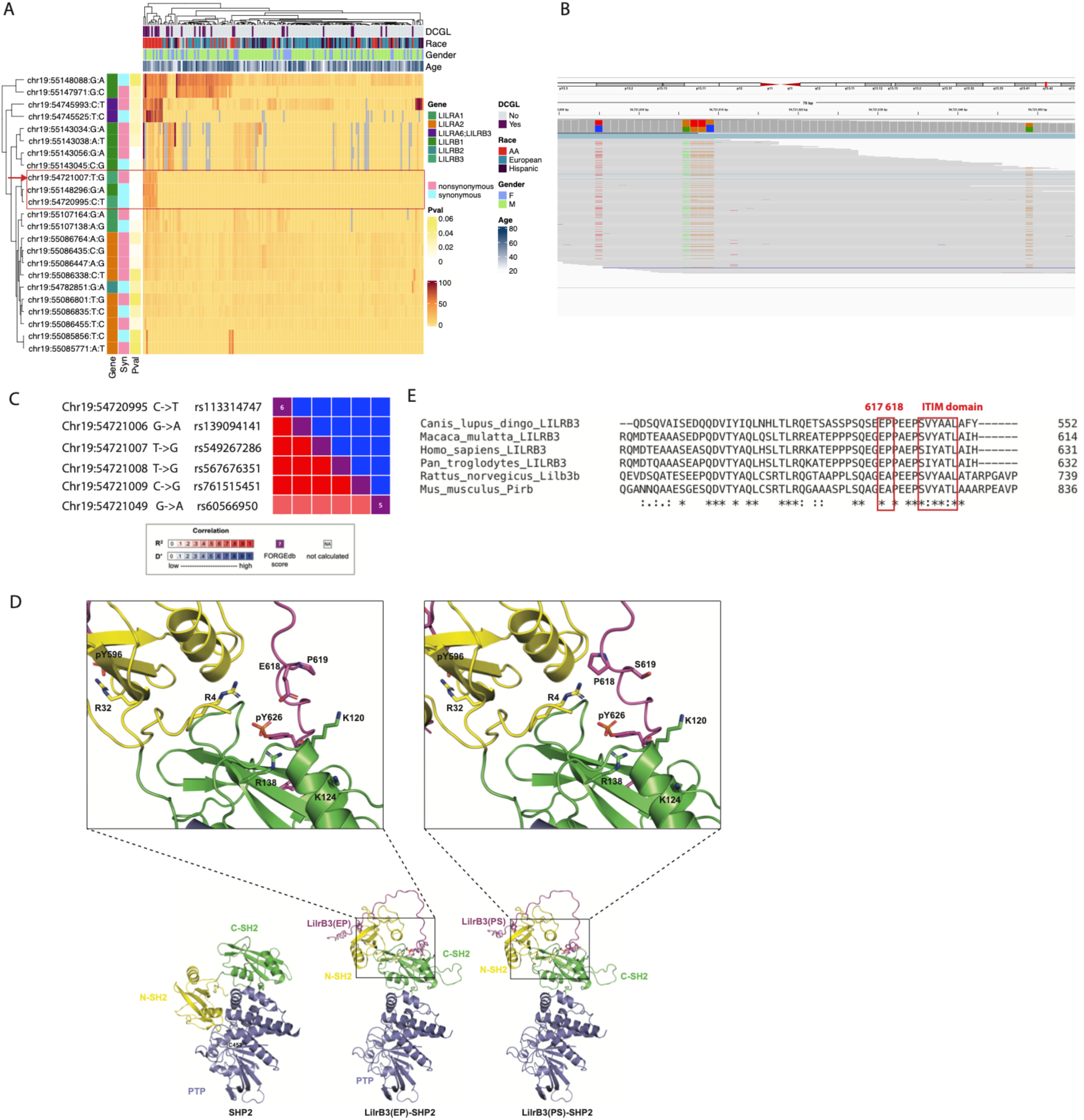
Identification of a cluster of 4 consecutive missense SNPs in *LILRB3* gene (*LILRB3*-4SNPs) associated with post-transplant renal failure. A) Two-way cluster view of LILRB eSNPs by AEF and demographics/outcomes; B) Sequencing alignment of reads covering 4 consecutive SNPs and their adjacent SNPs within *LILRB3* by IGV (chr19:54720990-54721060, hg19); C) Heatmap of the R^2^ and D’ between the *LILRB3*-4SNPs and their 2 adjacent SNPs; D) Protein structure modeling of the interaction of SHP-2 with *LILRB3* harboring *LILRB3*-4SNPs or reference alleles. E) Protein sequence alignment of *LILRB3* orthologs across organisms, human (Homo sapiens), chimpanzee (Pan troglodytes), monkey (Macaca mulatta), dog (Canis lupus dingo), rat (Rattus norvegicus) and mouse (Mus musculus).

When we stratified patients into three groups based on the AEF of *LILRB3*-4SNPs: High (AEF >= mean AEF), Low (0 < AEF < mean AEF), and no expression (AEF = 0), sequence alignment showed that the expression of this SNP (AEF > 0) was predominant in the non-European population (AA + Hispanic), with a notably much higher frequency within the AA population (**Figure 4A**). Patients demonstrating relatively elevated expression of the alternative allele of *LILRB3*-4SNPs exhibited a significantly heightened risk of graft loss within the entire RNAseq cohort (n=170, KM log-rank p < 0.0001, **Figure 4B**), within non-European recipients (n = 75, KM log-rank p = 0.0019), and within AA recipients (n = 40, KM log-rank p=0.002, **Figure 4C** and **Figure S3A**). The overall expression of *LILRB3* was not associated with graft loss (**Figure S3B**), suggesting that the *LILRB3*-4SNPs contributes to graft loss through the modulation of *LILRB3* function rather than the regulation of gene expression. To scrutinize whether the genotype of this variant correlates with graft loss at the DNA level within the non-European patients, targeted DNA sequencing of this SNP locus was undertaken on blood samples from 127 non-European individuals within the GoCAR cohort (**Methods**). Analyzing the subset of 32 individuals with both RNA and targeted DNA sequencing data, we initially verified the genotyping of the variant from both technologies. Importantly, we discovered a significant association between the *LILRB3*-4SNPs and DCGL within the 127 non-European individuals (**Figure S3C**), as well as within AA samples (**Figure 4D**). As anticipated, the genotype data from the combined RNA and targeted DNA sequencing cohorts demonstrated a notable association with graft loss in AA (n = 83) (**Figure 4E**) or AA+Hispanic (n = 169) population (**Figure S3D**). Among 83 AA recipients, 16 (19.3%) patients possessed LILR3-4SNPs, of whom 11 (68.8%) lost their grafts within 5 years post-transplant. No significant difference of other demographic and baseline clinical information (HLA-mismatch, donor status and induction therapy etc.), was found between the AA patients with and without *LILRB3*-4SPNs risk allele (**Table S5**). *APOL1* exonic variants (G1, G2) also bear relevance to the AA-recipients [16-19]. We assessed the co-occurrence of *APOL1* variant alleles with *LILRB3*-4SNPs and found that *APOL1* variant and *LILRB3*-4SNPs were not correlated by expression (**Figure S4)** or genetic linkage (**Table S6)** in our cohorts. After adjusting by *APOL1* genotype (one risk allele and two risk alleles) *LILRB3*-4SNPs remains significant with DCGL (**Table S7**). By cox regression analysis, LILR3-4SNPs is associated with graft loss (hazard ratio (HR)=3.58, p=0.002) adjusted for donor status and HLA mismatches (**Table S8**). The logistic regression test showed a highly significant association between SNP the graft loss within 5 years p=0.003 and odd ratio=7.38), together indicating that *LILRB3*-4SNPs is a distinct polymorphism enriched in AA transplant recipients and has a strong association with transplant graft failure.

**Figure 4.**
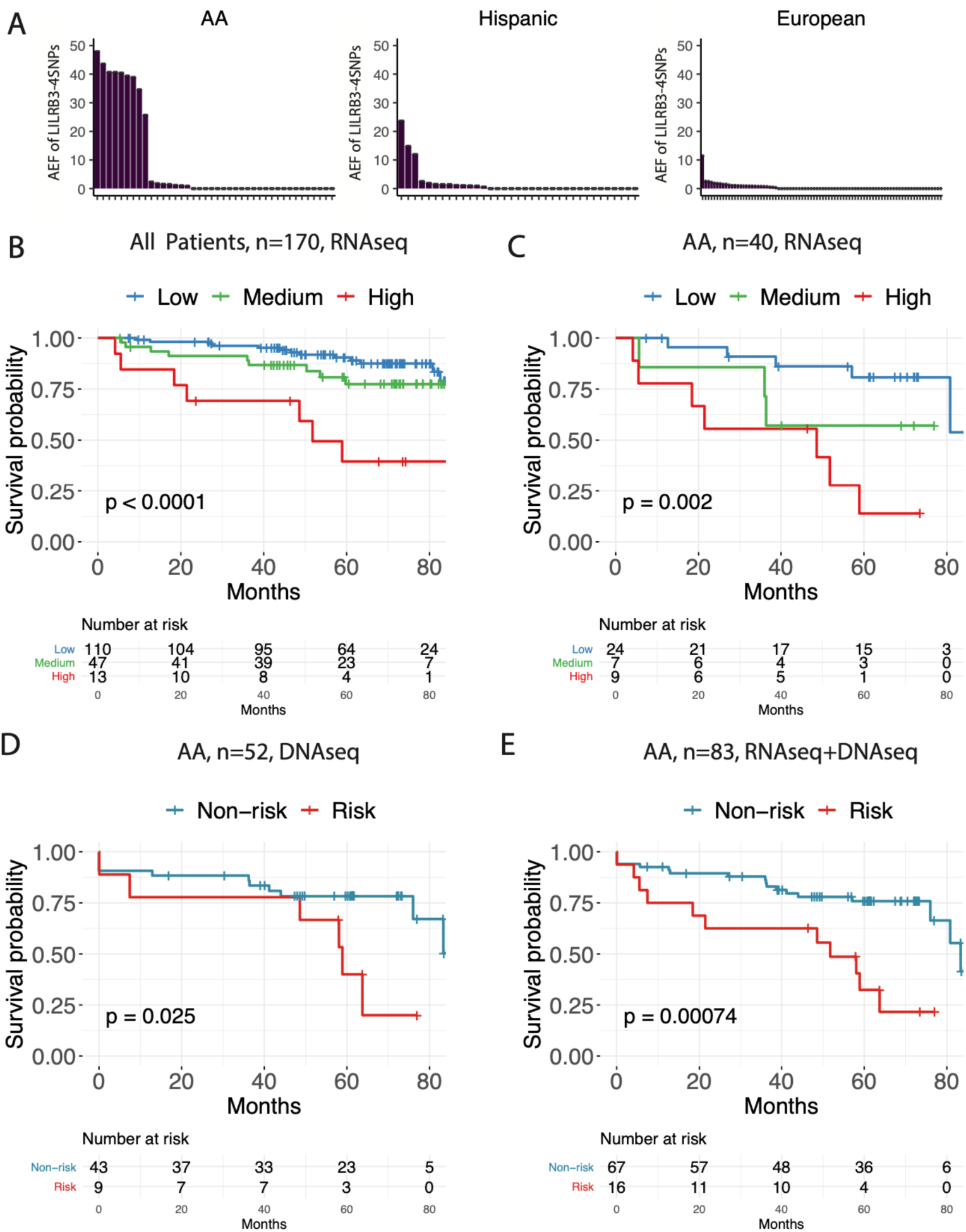
The association of *LILRB3*-4SNPs with post-transplant renal failure in GoCAR cohort. **A)** AEF of the *LILRB3*-4SNPs within AA (n = 40), Hispanic (n = 37) and European (n = 83) patients in GoCAR pre-transplant RNAseq cohort. The association of *LILRB3*-4SNPs with graft loss in the entire cohort (n = 170, **B**) or only AA population (n = 40, **C**) by RNA sequencing and validation by DNA sequencing (n = 52, **D**). The association of the genotype of *LILRB3*-4SNPs with graft loss in RNA+DNA sequencing cohort (n = 83, **E**).

### Genetic differentiation of the *LILRB3*-4SNPs and signatures of natural selection

We examined differentiation of *LILRB3*-4SNPs both at the RNA and DNA levels, employing three RNA-seq datasets each containing multiple genetic- or self-reported racial groups (CTOT19, VericiDx, and whole exome sequencing (WES) data from the BioMe biobank, along with the 1000 Genomes database (**Table S9**). The expression of the variant allele was specifically detected in the whole blood of non-Caucasian populations, predominantly observed within the AA samples across independent RNA sequencing datasets. In our extensive WES analysis of the substantial BioMe dataset (n=30,099), *LILRB3*-4SNPs predominantly manifested within the AA samples (n=7096, minor allele frequency (MAF) of 4.66%, presenting in 608 patients (8.57%)) and rarely in the Hispanic population (1.57% MAF or 2.3% prevalence) and while the European displayed negligible frequencies (<0.1% MAF or 0.1% prevalence). This distribution pattern aligns with the observations from the dbSNP database (MAF: 4.92% in AA, 0.9% in Hispanic and 0% in European) (**Figure S5**) ^17^.

To explore whether this racial discrepancy could be attributed to evolutionary positive selection within the LILR region, we scrutinized the rank scores of positive selections across diverse population samples, utilizing multiple population genetic summary statistics calculated for the CEU (Northern Europeans from Utah), CHB (Han Chinese in Beijing) and YRI (Yoruba population in African) samples from the 1000 Genomes Selection Browser. **Figure S6** shows the values of various summary statistics that can highlight patterns of genetic diversity consistent with recent selective sweeps. Pairwise *F*_ST_ and XP-CLR^18^ value comparisons involving the CHB population sample reveal a plateau of elevated values in LILR locus, consistent with previous evidence of recent positive selection found in Asian population samples^19^. We also observed elevated values of Fay and Wu’s *H* in CHB^20^, and to a lesser extent in CEU and YRI. We furthermore detected a small peak value of SweepFinder’s CLR test statistic^21^ in YRI. Together, these results suggest that recent positive selection in the LILR region may not have been limited to Asia. While we cannot be certain of the precise target(s) of selection in this region, these data do suggest that positive selection has acted on the LILR region in recent human evolutionary history. In addition, a recent study searching for signatures of positive selection acting on gene expression levels found evidence of selection in LILR genes ^22^. These findings are consistent with the notion that genes with immune-related functions often participate in evolutionary arms races with pathogens^23^, with MHC genes (which interact with LILR genes) showing especially strong evidence of recent selection ^24,25^.

### *LILRB3*-4SNPs associates with transcriptomic evidence of enhanced proinflammatory immune response and monocyte ferroptosis in pretransplant blood

To access the potential functional effects of these SNPs, we undertook meta-differential analysis of three RNAseq datasets from pretransplant blood in AA recipients (GoCAR, CTOT19, and VericiDx) and identified a total of 1755 meta-differentially expressed genes (mDEGs) associated with the *LILRB3*-4SNPs risk allele (**Table S10** and **Methods**). Among these genes, there were 892 upregulated and 663 downregulated (P < 0.05) (**Figure 5A)**, with a particularly interesting observation: Ferritin Light chain (*FTL*) emerged as the most downregulated gene (**Figure 5B**). The upregulated genes displayed a strong connection to functions or pathways that include T cell activation, chromatin organization, whereas downregulated genes were linked to functions including erythrocyte/myeloid cell differentiation, oxygen transport, oxidative phosphorylation, and mitochondrial function signatures (**Figure 5C**). Exploring the cellular specificity, upregulated genes were enriched with B/T cell-specific signatures, while downregulated genes were characterized by monocyte-specific signatures (**Figure S7**), which was consistent with that for differentially expressed serum proteins identified by mass spectrometry in the recipients carrying SNPs (n = 8) compared to those without the SNPs (n = 8) (**Figure S8, Table S11** and **Methods**).

**Figure 5.**
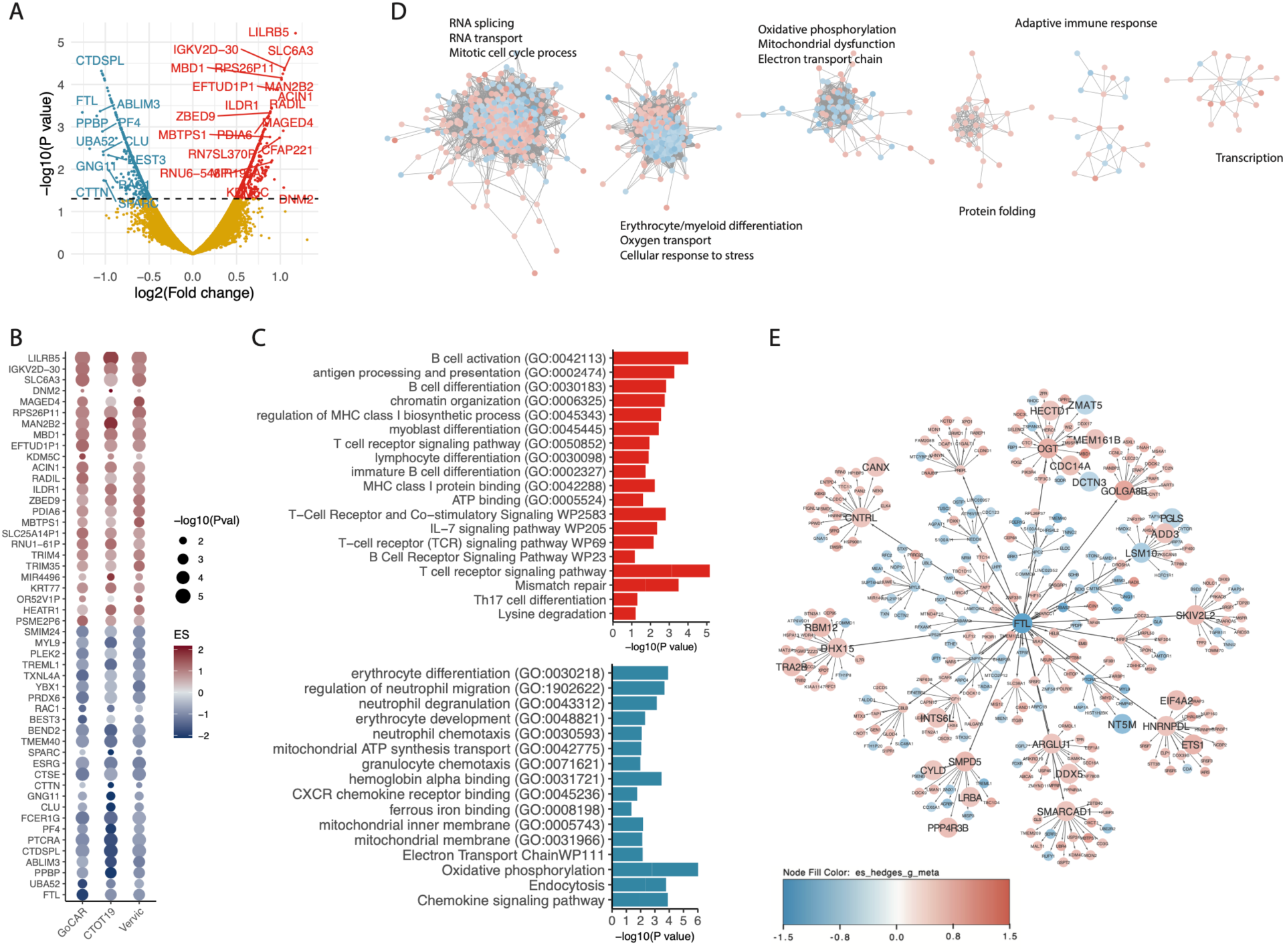
Differentially expressed genes (DEGs) associated with *LILRB3*-4SNPs by meta-analysis of 3 pre-transplant blood RNAseq datasets (GoCAR, CTOT19 and VericiDx). A) Volcano plot of meta DEGs between patients with and without *LILRB3*-4SNPs in 3 cohorts; **B**) Dotplot of top 20 up- and down-regulated meta genes across cohorts; **C**) the functional categories associated with meta DEGs; **D**) An overview of top 5 largest meta-clusters. **E**) A simplified network of key driver gene FTL, demonstrating the closest 2 layers of connections in its downstream. Enlarged nodes indicate key drivers, with FTL in the center is a global driver and others are local drivers.

We further performed a meta-correlation network analysis on the mDEGs in patients carrying the *LILRB3*-4SNPs across the three RNAseq datasets ^26^. This approach allowed us to construct a meta-correlation network, segmented into various sub-modules (**Figure 5D**). To ascertain the driving factors within modules housing at least 10 genes, we applied a driver analysis^26^. Notably, among these modules, the largest submodule containing 679 genes was notably driven by *FTL* (**Figure 5E**). Given that *FTL* functions as a model for iron storage, the reduced expression of *FTL* can elevate cellular iron levels, consequently fostering ROS accumulation and culminating in cell death by ferroptosis^27^. These analyses suggest that *LILRB3*-4SNPs may enhance immune responses (as evidenced by increased TNFα levels), subsequently leading to the downregulation of *FTL* ^28^, that could in turn contribute to immune mediated graft loss.

To delve into the extent of transcriptional dysregulation linked with the *LILRB3*-4SNPs within distinct immune subsets of kidney transplant patients, we embarked on a single-cell RNA sequencing analysis using pretransplant PBMCs derived from two African American transplant recipients with and without *LILRB3*-4SNPs (with matched demographic information). We observed larger T cell population and smaller monocyte population in the patients carrying the SNPs (**Figure 6A and B**). Among the discernible immune subsets, we notably detected a predominant expression of *LILRB3* within the CD16+ and CD14+ monocyte populations (**Figure 6C**). This insight directed our focus towards conducting a transcriptomic assessment specifically on monocytes. We carried out a differential expression analysis on monocytes between the two patients, resulting in 1076 differentially expressed genes (491 upregulated and 585 downregulated) within the CD16+ monocytes (**Figure S9A**). We observed a significant overlap (p < 0.001) between these differentially expressed genes in monocytes and those identified across the whole blood RNAseq profiles (**Table S12**). *FTL*, one of the top downregulated DEGs in monocytes, was also detected within this shared set of genes. The upregulated differentially expressed genes appeared to be involved in functions and pathways associated with type I/II interferon signaling, T cell receptor and co-stimulation signaling, and antigen presentation. Conversely, the downregulated genes were predominantly tied to functions related to mitochondria ATP synthesis and transport, oxidative phosphorylation, and remarkably, ferroptosis (**Figure 6D**). Similar results were observed in CD14+ monocytes (**Figure S9 B** and **C**). Among the ferroptosis-negatively-associated DEGs (e.g., *FTL, FTH1, GPX4, TFRC*) expressed prominently within the CD16+ and CD14+ monocyte subsets (**Figure 6E**), exhibited a reduced expression pattern in the monocytes of patients with the *LILRB3*-4SNPs (**Figure 6F**). This observation is consistent with earlier reports that indicated a diminished expression of these genes correlating with the onset of ferroptosis ^27^. Collectively, our single-cell RNA sequencing analysis of PBMCs implicated that the *LILRB3*-4SNPs cluster contributes to an enhanced immune response, while concurrently inducing ferroptosis primarily within the monocytes of PBMCs.

**Figure 6.**
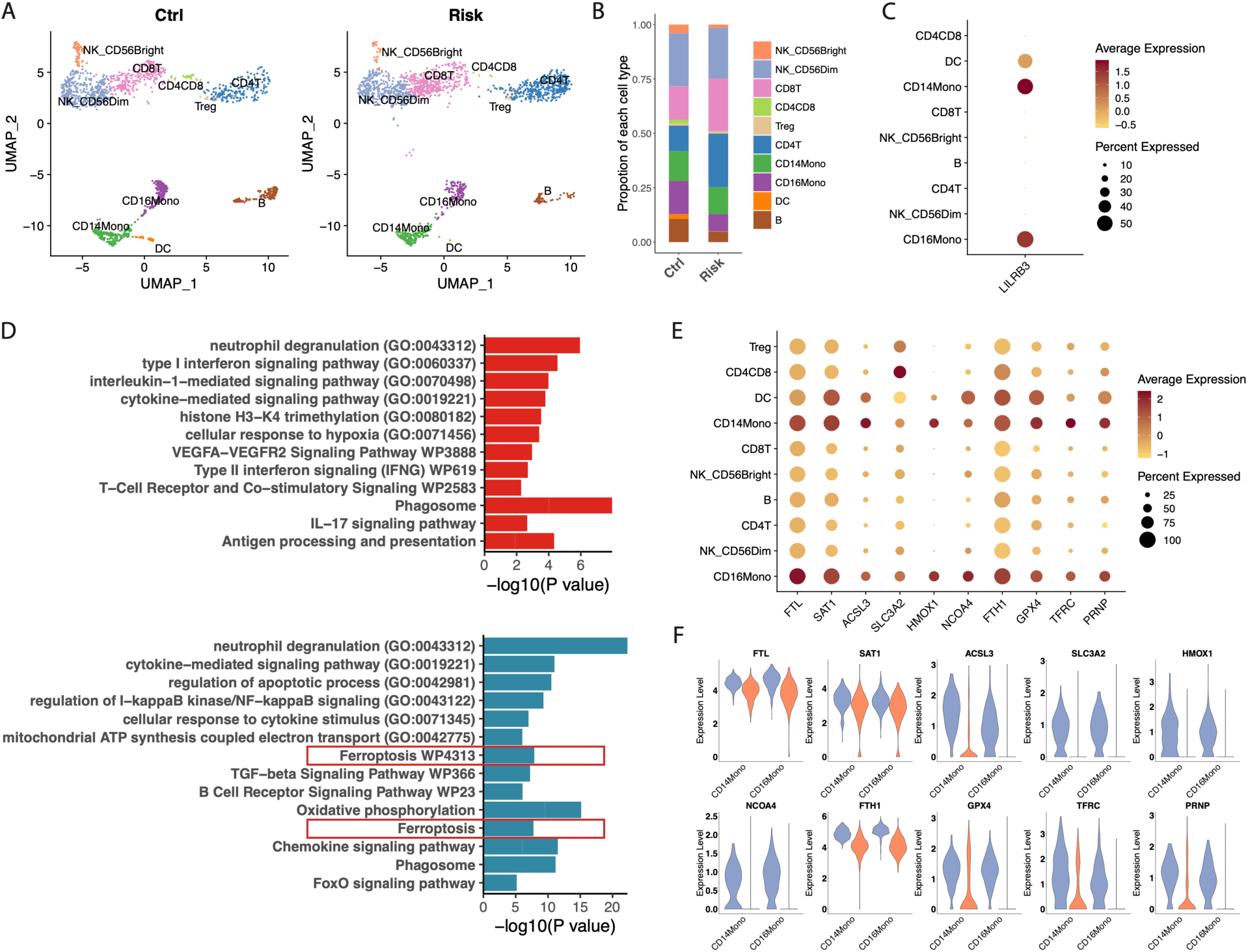
Single cell RNAseq of the pre-transplant PBMC from 2 AA recipients with and without *LILRB3*-4SNPs risk allele. **A)** UMAP showing the cell population of 2 PMBC samples. **B**) Cell proportion of each cell type within each sample. **C**) The expression of *LILRB3* in each cell type. **D**) Function enrichment of genes significantly up- and down-regulated in patients with risk allele within CD16+ monocytes. **E**) Expression of ferroptosis-related DEGs in each cell type. **F**) Expression of ferroptosis-related DEGs between risk and non-risk sample within CD14+ and CD16+ monocytes.

### *LILRB3*-4SNPs associates with transcriptomic evidence of persistent inflammation and monocyte ferroptosis in post-transplant blood and kidney allograft biopsies

Most of AA recipients received T cell depleting, rabbit anti-thymocyte globulin (rATG) induction therapy at the transplantation. Previous studies by our collaborators among others ^29^, show immune recovery by 6-12-month posttransplant. RNA sequencing of the whole blood collected after 6-months post transplantation from 20 AA recipients with (n = 10) and without (n = 10) *LILRB3*-4SNPs showed the upregulation of T cell receptor signaling and B cell mediated immunity (**Figure 7A**). Single cell RNA sequencing on the PBMC isolated from the two AA recipients (one with and one without *LILRB3*-4SNPs) with matched demographic and clinical features (age, gender, HLA mismatches, and ATG induction) at 24 months post-transplant was performed. Consistent with the findings in pre-transplant peripheral blood, we observed larger T cell and smaller monocyte populations in the SNPs-carrying recipient (**Figure 7B and C**). The DEG analysis exhibited similar dysregulated pathways comparing to pretransplant blood including of upregulation of T/B cell activation and down-regulation of ferroptosis-associated genes in monocytes (**Figure 7D and E**, and **Table S13**).

**Figure 7.**
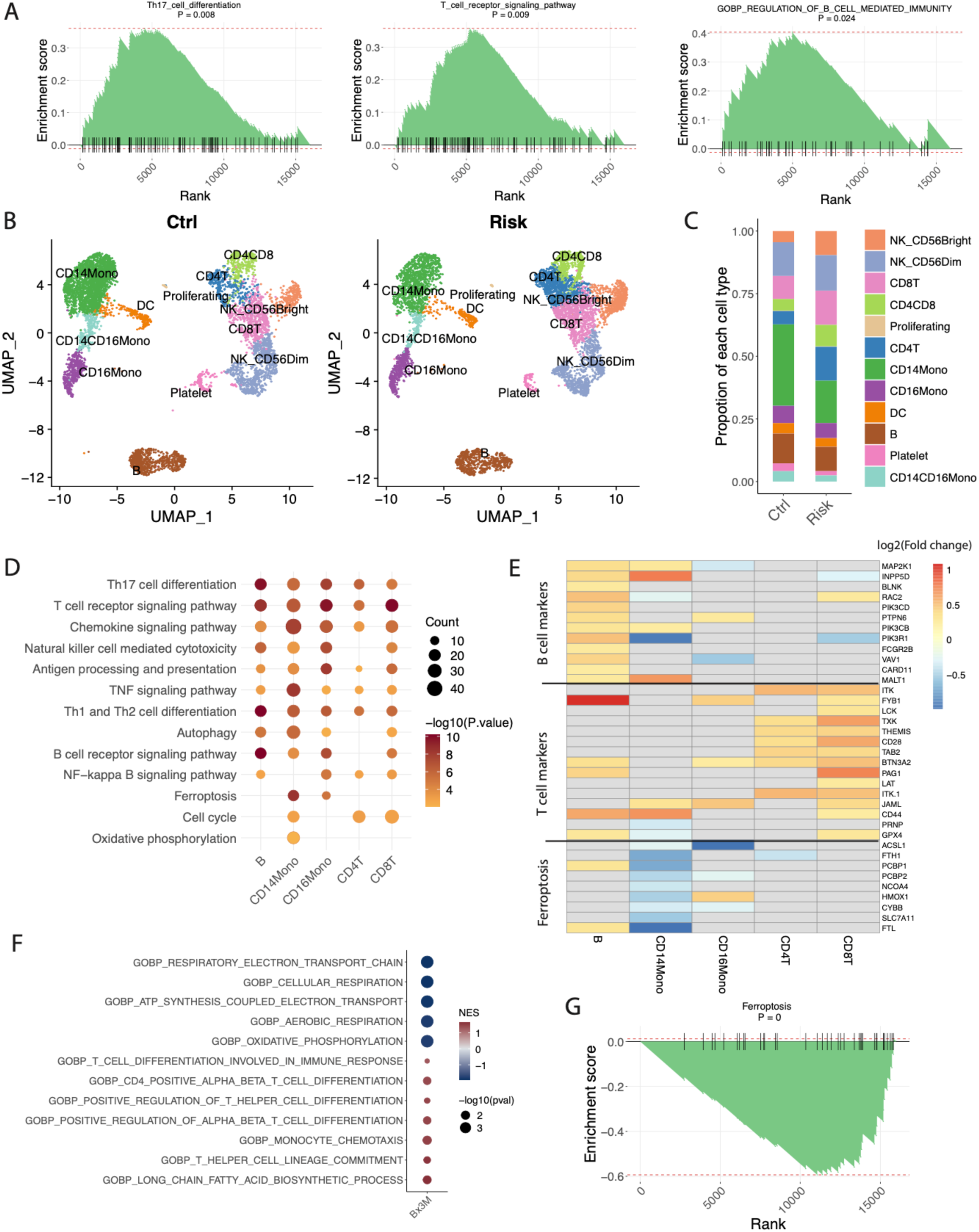
Transcriptomic dysregulation of post-transplant blood and kidneys in AA recipients carrying *LILRB3*-4SNPs. **A)** GSEA enrichment plot of the pathways of Th17 cell differentiation, T cell receptor signaling and B cell mediated immunity in bulk RNA sequencing of the blood samples collected after 6 months post-transplant in 20 AA recipients with (n = 10) and without (n = 10) *LILRB3*-4SNPs. **B**) UMAP of the cell population in the PBMCs isolated from 2 AA patients (with and without *LILRB3*-4SNPs risk allele) at 24-month after transplantation. **C**) Cell proportion of each cell type in each patient. **D**) Function enrichment of significant DEGs between patients with and without *LILRB3*-4SNPs in each cell type. **E**) Heatmap showing the log2(fold change) of selected DEGs in each cell type. **F**) Significantly-dysregulated functions (GSEA enrichment score) in 3-month post-transplant biopsies of 6 AA recipients with (n = 3) and without (n = 3) *LILRB3*-4SNPs risk allele. **G**) GSEA enrichment plot of Ferroptosis pathway comparing the patient with (n = 3) and without (n = 3) *LILRB3*-4SNPs risk allele in 3-month biopsies.

We next investigated the association of the recipient’s *LILRB3*-4SNPs with the gene expression dysregulation in post-transplant kidneys. We identified 3 pairs of AA kidney transplant recipients with and without the *LILRB3*-4SNPS with matched donor and recipient’s demographic and clinical parameters from the published expression dataset from GoCAR cohort ^7^ and performed a paired test to identify DEGs in biopsies associated with the recipient’s *LILRB3*-4SNPs (**Table S14**). Similar to pre- and post-transplant blood profiles, we observed the over-expression of genes involved in T/B cell activation and down regulation of genes in oxidative phosphorylation, mitochondria electron transport (**Figure 7F**) in the post-transplant kidneys in the recipients with *LILRB3*-4SNPs. By GSEA analysis, we also detected the down-regulation of ferroptosis-negatively-associated gene set that were seen in the pretransplant blood (**Figure 7G**). The shared transcriptional dysregulation between recipient’s pretransplant blood and post-transplant kidneys implied the persistent inflammation in the blood stream post-transplant causes kidney damage.

### *LILRB3*-4SNPs associates with post-transplant worse graft function, pathology and complications

We next tested for associations of graft function, pathological lesions and complications with LRB3-4SNPs in the GoCAR, CTOT19 and kidney transplant patients in the BioMe cohort. The recipients with the SNPs exhibited declined eGFR (>30% decrease from 6-24 month or consistently below 30) in the GoCAR cohort (p=0.017, **Figure S10**), accelerated eGFR declining in BioMe (**Figure S11**) cohort, IFTA Grade II or eGFR declining (>2, p=0.01) in the CTOT19 cohort (**Table S15**). The *LILRB3*-4SNPs was associated with the high incidence of glomerulitis (Banff g>1) in the GoCAR (p=0.044, **Table S16**) and BioMe (p=0.004, **Table S17**) cohorts. Glomerulitis is one key factor for chronic transplant glomerulopathy and renal failure ^30^ and also associated with graft loss in GoCAR cohort (g at 3 m, p=0.04; g at12 m, p=0.04; g at 24m, p = 0.01) (**Table S18)**. Interestingly, In BioMe cohort, *LILRB3*-4SNPs was also linked to the development of sepsis in kidney transplant patients (p=0.010) (**Table S19**). Sepsis, a common complication in kidney transplant and ESRD, one of the leading causes of kidney failure and mortality ^31–38^.

### *LILRB3*-4SNPs induced inflammation and ferroptosis, which was attenuated by a ferroptosis inhibitor *in vitro*

We conducted a series of *in vitro* functional assays to delve into the causal association of *LILRB3*-4SNPs with enhanced inflammation and ferroptosis in monocytes. Pretransplant PBMCs extracted from 6 AA kidney transplant recipients (3 with and 3 without SNPs) were treated with *LILRB3* agonists or antagonists while subjected to 6-hour LPS stimulation. The PBMCs carrying the *LILRB3*-4SNPs (**Figure 8A, left,** one representative patient,) exhibited a heightened release of TNFα compared to their counterparts lacking the SNPs (**Figure 8A, middle,** one representative recipient). Notably, the agonist (**middle bar**) exhibited a less pronounced inhibitory effect on PBMCs with the *LILRB3*-4SNPs in comparison to the recipients without the SNPs (**Figure 8A, middle)** or healthy donor PBMCs (HD208) (**Figure 8A, right)** implicating a disrupted inhibitory response of *LILRB3* caused by the SNPs.

**Figure 8.**
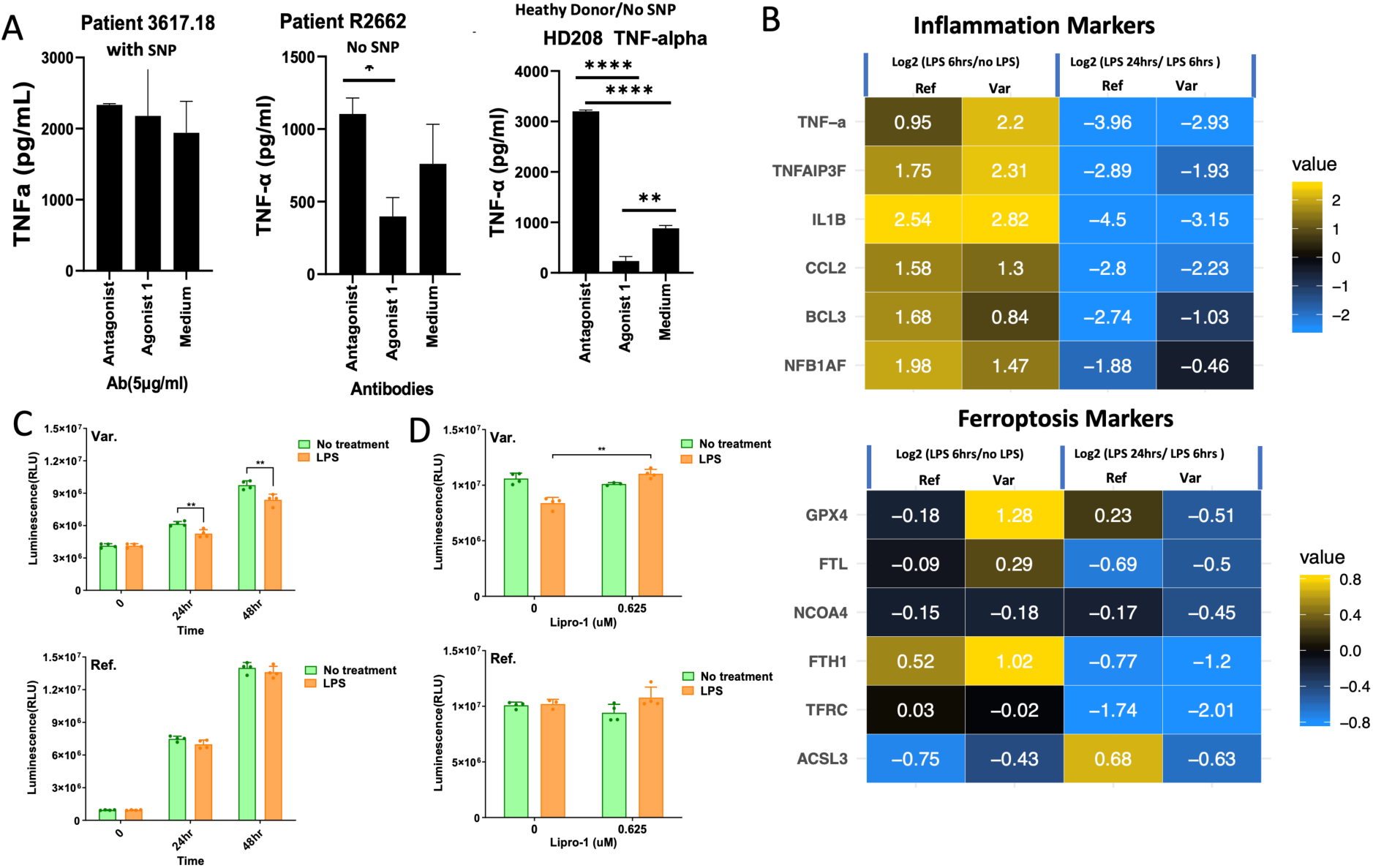
*In vitro* functional analysis of *LILRB3*-4SNPs. **A)** TNFa release assay in PBMCs upon *LILRB3* antagonist/agonist treatment; qPCR on immune response and ferroptosis-negatively-associated genes **(B)**, survival analysis within 48 hours upon LPS stimulation **(C)** and in conjunction with ferroptosis-inhibitor (lipro-1) treatment **(D)** in THP-1 cells overexpressing *LILRB3*-4SNPs and reference alleles.

To further test the effects of *LILRB3*-4SNPs on immune cell function we generated two lentiviral constructs overexpressing *LILRB3* gene carrying the alternative (Var) or reference (Ref) allele of the *LILRB3*-4SNPs. These constructs were then transduced introduced into THP-1 (macrophage) cell lines. Consistent with a previous report ^14^, following a 6 h LPS stimulation, all cell lines exhibited increased expression of crucial inflammatory response markers affiliated with the LILRB family, including TNFα and IL1β, and the expression of these genes decreased from 6 to 24 hours (**Figure 8B, upper**). The cell line overexpressing the Var allele produced greater quantities of TNFα, TNFAIP3 and IL1β upon 6-hour LPS treatment, with less attenuation between 6 and 24 hours, implicating increased inflammation associated with the SNPs (**Figure 8B, upper**). Expression of 5/6 ferroptosis negatively-associated genes in cell line with the SNPs decreased more between 6 and 24 hours post LPS treatment, consistent with enhanced ferroptosis at 24 hours linked to the SNPs (**Figure 8B, lower and right**).

Survival analysis demonstrated a reduced viability upon LPS stimulation in cells with the SNPs (**Figure 8C, upper,** orange vs green) but not those without the SNPs (**Figure 8C, lower,** orange vs green). This phenomenon for the SNPs was reversed by ferroptosis inhibitor, Liproxstatin-1 (Lipro-1, at 0.625 umol) that targets lipid peroxidation (orange bar in **Figure 8D, upper)**. Lipor-1 had no effect on the cells without SNPs (orange bar in **Figure 8D, lower**). These findings using SNP overexpression collectively implicated that the *LILRB3*-4SNPs activates the NFKB pathway which in turn induces the expression/release of TNFα and cytokines and chemokines linked to LILRB3 (such as IL1B, CCL20), and subsequently, this cascade appears to suppress the expression of ferroptosis-negatively-associated genes (eg. ferritin and GPX4), ultimately culminating in ferroptosis within monocytes.

### *LILRB3*-4SNPs associates with the severity or complications in kidney and other immune-related diseases in a large EHR-linked biobank cohort (BioMe)

As LILRB families plays a central role in regulating immune responses [12], we hypothesized that, in addition to its involvement in the adverse outcomes of kidney transplantation, *LILRB3*-4SNPs might also be associated with immune-related conditions in the broader African American (AA) population. To explore this, we tested for relationships among *LILRB3*-4SNPs and various health conditions (defined by ICD10 code) in a cohort of 7,096 AA patients with electronic health records in the BioMe database (**Methods**). While we did not observe an association between these SNPs and the occurrence of kidney diseases, our analyses showed that these SNPs were linked to the occurrence of other health issues, including respiratory diseases, pain, and viral/bacterial infections (**Table S20**), associations that are distinct from the primary association of *APOL1* G1/G2 alleles with kidney diseases (**Table S21**)^39–41^.

Based on this data, we hypothesized that the link between *LILRB3*-4SNPs and kidney diseases might impact on disease progression rather than disease onset. To explore this, we first evaluated the historical eGFR value in each patient and estimated the age of first eGFR decline (≤ 15) (**Methods**). Survival analysis suggested ESRD patients with *LILRB3*-4SNPs have significant early decline comparing to those without the *LILRB3*-4SNPs risk alleles in AA population (**Figure 9A and B**) while patients with 2 *APOL1* risk alleles showed worse outcome in both ESRD and CKD (**Figure 9C and D**) as described in previous studies^42,43^. After adjusting *APOL1* genotype, *LILRB3*-4SNPs risk alleles remain significantly associated with early eGFR decline in ESRD patients, independent of *APOL1* risk alleles (**Table S22**). By further grouping patients with *APOL1* 2 risk alleles alone, *LILRB3*-4SNPs alone or combined (with both *APOL1* 2 risk alleles and *LILRB3*-4SNPs risk alleles), the combined group showed higher hazard ratio, suggesting additive effects of both genotypes (**Figure 9E-F** and **Table S23**). The significant association of *LILRB3*-4SNPs risk allele alone with early eGFR decline in ESRD patients, further confirming the independent association of *LILRB3*-4SNPs with ESRD progression (**Figure 9E**, **Table S23**).

**Figure 9.**
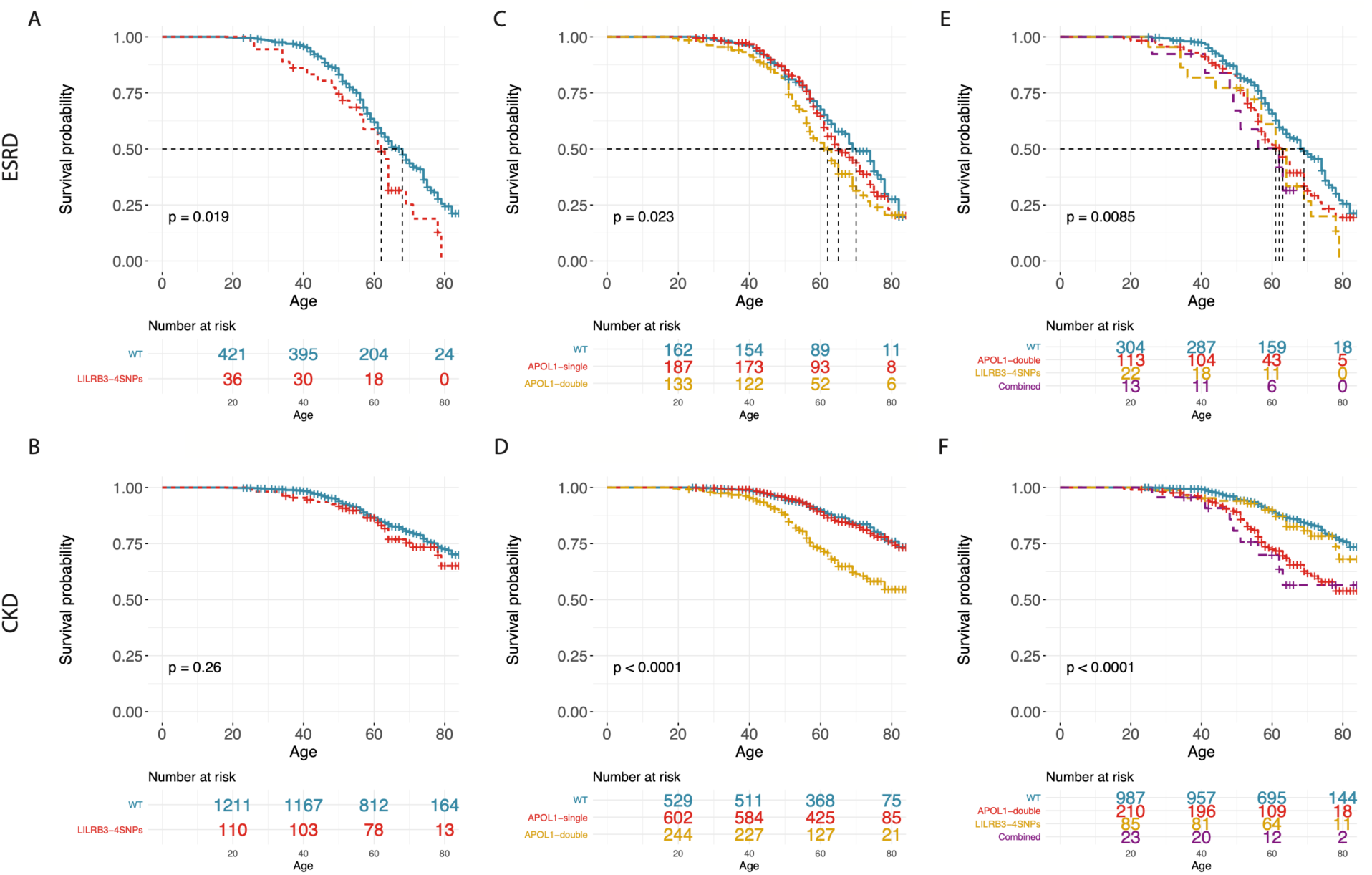
Survival analysis of *LILRB3*-4SNPs or *APOL1* G1/G2 alleles with the eGFR decline (<= 15) in the AA patients with kidney diseases in BioMe cohort. Survival curve (p value by log ratio test) of the eGFR decline (<= 15) of, ERSD and CKD patients with AA ancestry in Biome Biobank grouped by *LILRB3*-4SNPs alone (**A and B**), APOL1 single or double alleles (G1/G1, G2/G2 or G1/G2) alone (**C and D**) or combined alleles (*LILRB3*-4SNPs+*APOL1* double alleles) (**E and F**).

Subsequently, we examined the correlation between *LILRB3*-4SNPs and complications linked to kidney diseases. We identified a notable correlation between *LILRB3*-4SNPs and sepsis in patients with end-stage renal disease (ESRD) (p=0.008) and chronic kidney disease (CKD) (p=0.024), consistent with the findings observed in kidney transplant patients as described earlier (**Table S19**). In contrast, the G1/G2 double alleles of *APOL1* exhibited a weaker correlation with this condition when compared to *LILRB3*-4SNPs (**Table S19**).

Collectively, our observations within the large BioMe cohort indicate that *LILRB3* is linked to the progression and complications of kidney diseases (notably ESRD), independent of *APOL1* G1/G2 allele. The combination of *LILRB3*-4SNPs and *APOL1* G1/G2 alleles exhibited an additive effect on the CKD/ESRD progression. Moreover, in comparison to the *APOL1* G1/G2 allele, *LILRB3*-4SNPs exert a broader influence on immune-related disorders beyond kidney transplantation and diseases prevalent within the African American population.

## Discussion

Genetic factors may contribute to a high risk of renal failure in African American kidney transplant recipients. Using RNA sequencing along with targeted sequencing and whole exome sequencing in the cohorts from multiple studies, we for the first time identified the AA-specific *LILRB3*-4SNPs as a strong risk factor for kidney transplant failure and it also has a broad association with the severity of kidney and other immune related diseases in African American patients.

*LILRB3* as one member of LILR gene family is primarily expressed in myeloid cells. It functions as a negative immune response regulator by binding to SHP-1/2 through ITIM motif to suppress the NFKB pathway, resulting in reduced cytokine release and inflammation^14^. The ligand for *LILRB3* is not clear although its upregulation in synovial tissue was associated with RA and elevated expression was detected in breast and colon cancers^13^. Although other LILR gene family members (e.g. *LILRB1* and *LILRB2*) were known to mediate transplant tolerance through binding to HLA-G ^13^, the role of *LILRB3* in kidney transplant or diseases is largely unknown. Here we demonstrated the missense polymorphisms in the close proximity to the ITIM motif but not the total expression of *LILRB3* gene in peripheral blood had the effect on the transplant outcomes. The SNPs enhanced the adaptive immune responses and ferroptosis in monocyte in pre- and post-transplant peripheral blood, resulting the graft damages in kidney. The molecular and cellular mechanism by which *LILRB3*-4SNPs regulates the immune response in peripheral blood and subsequently impacts on the graft remains to be extensively investigated. Firstly, the binding capability of *LILRB3* with the SNPs to SHP-1/2 proteins can be examined by *in vitro* biophysical binding assay or crystallography. Secondly, the immune response activity or cell survival in immune subsets and their interactions associated with the SNPs in peripheral blood can be investigated in depth such as using *in vitro* co-culture of LPS-stimulated monocytes with T or B cells isolated from the recipients carrying the SNPs. In addition, in the graft, the lymphocyte infiltration and associated pathological lesions in the recipients carrying the SNPs can be investigated in the spatial context at the molecular level using spatial transcriptomics on the kidney biopsy samples. Lastly, we performed functional assays on THP-1 cell line over-expressing *LILRB3*-4SNPs to establish the causal role of the SNPs in the enhanced inflammation. Due to limitation of overexpression system, the CRISPR knock in (KI) of the SNPs in the original genomic sites could be used to study the effect of the SNPs on immune response under the regulation of *LILRB3* promoter, which reflects real biology of *LILRB3*. The humanized mouse transplant model with CRSIPR KI of the SNPs can be used to test the systematic effect of the SNPs on immune response activity in peripheral blood and the subsequent impact on the graft injury.

Consistent with previous reports, our analysis indicated that *APOL1* G1/G2 alleles were primarily associated with the onset of the kidney diseases and the individuals with two *APOL1* risk variants (13% of the population) had a higher risk of nephropathy and more severe outcomes than the individual with only one variant allele. Conversely *LILRB3-*4SNPs is not associated with the onset of kidney diseases, but the patients carrying the *LILRB3*-4SNPs exhibited more severe outcomes/complications than the patients with two copies of *APOL1* G1/G2 alleles (eg. G1/G1, G1/G2 or G2/G2), particularly in the setting of ESRD and kidney transplant. This could be partially explained by distinct molecular functions of these two proteins. The *APOL1* is primarily expressed in epithelial cells of various tissue and is localized to podocytes, arteriolar endothelium and proximal tubular epithelium in kidney tissues. *APOL1* may play a role in lipid exchange and transport throughout the body. *APOL1* variants may be associated with enhanced opening of cation channels, resulting in endolysosomal and mitochondrial dysfunction and altered autophagy, endoplasmic reticulum stress. In contrast, *LILRB3* is exclusively expressed in immune cells (primarily in myeloid cells) and function as negative immune response regulator. *LILRB3*-4SNPs may induce systematic inflammation and macrophage/monocyte ferroptosis, leading to tissue damages in kidney damage or ESRD. This also explain why *LILRB3* has a broad effect on the immune related diseases (respiratory diseases and pains). The patients with both *APLO1* double alleles and *LILRB3*-4SNPs would have the worst manifest of nephropathy because of malfunctions in ion channel activity and enhanced inflammation. In summary, *LILRB3*-4SNPs alone might not initiate the nephropathy, but cause severe kidney damages in the individuals carrying *APOL1* double alleles due to enhanced systematic inflammation. The association of the SNPs with other immune related diseases warrants further investigations.

Ferroptosis, an iron-dependent form of non-apoptotic cell death, has emerged as a significant mechanism in the development of chronic kidney diseases ^44^. It presents a potential and novel therapeutic target for managing CKD ^45^. Despite this, our understanding of ferroptosis in the context of kidney transplants remains limited. The discovery of the association between *LILRB3*-4SNPs and ferroptosis in monocytes in the bloodstream and post-transplant kidneys is a compelling advancement. Notably, our findings demonstrate that a ferroptosis inhibitor effectively reduced cell death in THP1 cells overexpressing *LILRB3*-4SNPs, suggesting its potential utility in conjunction with ATG induction for treating recipients carrying these SNPs. However, it is crucial to evaluate the efficacy of the ferroptosis inhibitor in an *in vivo* mouse transplant model to validate its therapeutic potential.

In a summary, *LILRB3*-4SNPs represents a crucial genetic risk factor for the severity of kidney transplant outcomes and immune-related diseases within the African American population. Further research will focus on extensively investigating its molecular and cellular mechanisms involved in disease progression.

## Methods

### Detection of expressional SNPs (eSNPs) from RNA sequencing data

The pipeline of detection of expressional SNPs (eSNPs) based on SNP array and RNA sequencing data from pretransplant blood in GoCAR cohort was depicted in Figure S1. Briefly, the quality control, SNP calling and imputation was performed on SNP array data of blood RNA samples from the kidney transplant recipients as described in previous study ^11^. The SNP calling from RNAseq data of pretransplant blood samples from the recipients was performed with the GATK best practice ^46^ under GATK/4.2.0.0 and further described in **Supplementary Methods**. The SNP genotypes from SNP arrays and RNAseq were then combined and annotated with AnnoVar version “2018Apr16”. The exonic SNPs were subsequently extracted for further analysis. To quantify the RNA sequencing reads that are mapped to these exonic SNPs, the nucleotides of those exonic SNP locus on human genome (hg19) were masked as “N” to avoid mapping bias towards reference allele. RNA sequencing reads were then aligned to the masked human genome with STAR 2.6.1d and duplicate reads were identified by Picard 1.93. The reads mapped to reference and alternative alleles of each SNP were counted with ASEReadCounter. The alternative allele expression fraction (AEF) of each SNP was calculated as read count for alternative allele / (the total read count for reference and alternative allele). The reads covering target SNP site was extracted with samtools 1.9 and the coverage was visualized with IGV 2.8.2.

The pipeline was also applied to CTOT19, VericiDx and sclerosis RNA sequencing dataset to detect the allelic specific expression of LILR3-4SNPs identified in GoCAR cohort.

### Targeted DNA sequencing of the LILR3-4SNPs locus

The DNA samples from 127 non-Caucasian (AA+Hispanic) recipients in GoCAR cohort were retrieved from GoCAR specimen biobank and were subjected to DNA sequencing of targeted region around the LILR3-4SNPs locus (+/- 100bp) by following Illumina 16S sequencing protocol. The details of the library and sequencing were provided in Supplementary Methods.

To determine genotypes of the targeted region around the LILR3-4SNPs locus, the reference sequences of 200pb upstream and 200bp downstream rs549267286 was extracted from human genome reference (hg19) to build the targeted reference genome using bwa/0.7.15. The DNA sequences were aligned to the targeted reference with bwa mem with default parameters. SNPs within this region were called by bcftools/1.9 with mpileup by default parameters for each sample and finally merged by bcftools merge.

### Bulk RNA sequencing on pre-transplant and post-transplant blood and data processing

Bulk RNA sequencing on pre-transplant blood from the recipients in CTOT19 and VericiDx cohorts was performed by VericiDx Inc. Total RNA was isolated from the whole blood stored in PAXgene tubes and then subjected to standard RNA sequencing libraries generation by following manufactory protocol. The pre-transplant sequencing libraries were then sequenced on NextSeq sequencer at 75 bp read length and an average throughput of 12.5M reads per sample.

The sequencing libraries of post-transplant blood in GoCAR cohort were generated by stranded ployA selection kit and were then sequenced on NovaSeqXplus sequencer at 150 bp read length and an average throughput of 30M reads per sample at Cornell Genomics Core Facility.

The raw sequencing data from pre- and post-transplant blood were first aligned to human genome hg38 with STAR 2.5.3a ^47^ and gene expression counts in each sample was quantified with HTseq both with default parameters. Limma voom ^48^ was used to test the gene expression between risk and non-risk samples with raw gene expression counts. Genes with p value < 0.05was identified as differentially expressed (DEG).

### Structure modelling of the interactions of SHP-2 and *LILRB3*

To model the interaction between SHP2 and the C-terminal tail of *LILRB3*, the sequences of the two SHP domains of human SHP2 (residues 1-218) and the tail segment of *LILRB3* containing the last two ITIM motifs (residues 581-631) was used as the input for the AlphaFold 2 structural prediction in the ColabFold webserver ^15,16^. AlphaFold 2 successfully predicted the expected binding modes between the two ITIM motifs and the two SH2 domains in SHP2, respectively, despite the absence of phosphorylation of the tyrosine residues in the motifs. The phospho-groups were added by using Charmm-GUI and manually adjusted in Coot ^49,50^. The model was then aligned based on the second SH2 domain with the crystal structure of full-length SHP2 in the inactive state (PDB ID: 4NWF), where the first SH2 domain obstructs the active site of the catalytic domain. Combining the SH2 domains from the AlphaFold model with the catalytic domain from the crystal structure led to a composite model of full-length SHP2 bound to the C-terminal tail of *LILRB3*, in which the first SH2 domain is pulled away from the catalytic domain, indicating an open active conformation. The EP->PS variant of the *LILRB3* in complex with SHP2 was predicted in the same manner. The structure figures were rendered using PyMOL (The PyMOL Molecular Graphics System, Version 2.0 Schrödinger, LLC.).

### Meta differential gene expression and network analysis

Meta-analysis was performed across the three datasets (GoCAR, CTOT19 and VericiDx) using combined effect size (ES) approach to identify differential meta-gene signatures between samples with and without LILR-4SNP risk alleles. Meta-genes were selected if the following conditions were met: 1) genes were mapped to at least 2 out of 3 data sets. 2) p-value of the meta-ES < 0.05; 2) individual ES was in the same direction, either up or down, in all mapped data sets. These meta-genes (1705 genes) were further used for meta-network construction. Significant Meta-Coexpression-Network was identified by selecting connections with meta |r| >= 0.6. We constructed the network for samples with *LILRB3*-4SNPs risk alleles only and performed Markov Cluster Algorithm (inflation parameter=2) to identify sub-clusters from the network. To identify key driver genes within each sub-cluster, we performed z-normalization per sample for case samples in each data set and combined normalized expression data to construct directional Bayesian network. Key Drivers (KD) were then selected as genes that had large downstream numbers and out-degrees. If a KD is not in the downstream of any other KDs in the same module, it becomes a global driver otherwise it is a local driver. Detailed methods of meta-analysis were described in our previous publication ^26^.

### Single cell RNA sequencing on pre- and post-transplant PBMC

Single cell RNA sequencing was performed on PBMCs isolated from the pretransplant blood of 2 AA recipients (1 without and 1 without *LILRB3*-4SNPs) and post-transplant (24 months) blood of 2 AA recipients (1 without and 1 without *LILRB3*-4SNPs) with matched recipient and donor’s demographic and baseline clinical features (age, gender, donor status, HLA mismatches and induction types). The PBMC samples were retrieved from the liquid nitrogen tank and thawed for scRNAseq library generation. Briefly, the viable PBMC were loaded into chromium microfluidic chips and barcoded with a 10x Chromium Controller (10x Genomics). RNA from the barcoded cells was subjected to sequencing library generation with reagents from a Chromium Single Cell 3′ v.2 Reagent Kit (10x Genomics) according to the manufacturer’s protocol. The library was sequenced on Illumina sequencer (NovaSeq 6000) at a 150 bp paired-end sequence length and 400 million read throughput per library. The QC, clustering and DEG analysis was done with Seurat and was described in Supplementary Methods.

### TNFα immune response assay

PBMCs were isolated using LymphoPrep medium (07851, STEMCELL Technologies) from patient’s blood. Total PBMCs were incubated with anti-*LILRB3* agonist or antagonist (5 μg/ml) or IgG overnight and then PBMCs were stimulated with LPS (50 ng/ml) for 6 hours. The supernatants were collected, and TNFα production was determined by ELISA (eBioscience).

### Generation of the construct and cell line overexpressing the SNP reference and variance allele

The N-terminal myc-tagged *LILRB3* cDNAs with and without *LILRB3*-4SNPs were synthesized and cloned into a retroviral vector that also carries a blasticidin resistance gene under the control of IRES (internal ribosomal entry site). The *LILRB3* cDNA contains a human growth hormone signal peptide, myc-epitope tag, and the coding sequence of mature *LILRB3* protein.

### Cell viability assay

Cells were seeded in 96 well culture plates at a density of 1 × 104 cells/well and treated with 200ug/L LPS and 0.625umol liproxstatin-1. At the end of treatment, cell viability was determined using CellTiter-Glo Luminescence Cell viability Assay kit (Promega) by measuring luminescence (ATP level) following the manufacturer’s instructions. Medium without cells were used as background luminescence. Values were presented as an RLU.

### Quantitative PCR

Total RNA was extracted from THP-1 using TRIzol Reagent (Invitrogen) after incubated with 200ug/L LPS for 6 hours and 24 hours; 1000 ng total RNA was reverse transcribed to cDNA with the EasyScript Plus™ cDNA Synthesis kit (Lambda Biotech). Quantitative real-time PCR was performed with the 7500 Real-Time PCR System (Applied Biosystems). The mRNA level was normalized to GAPDH and expressed as fold change. The primer sets can be found in the supplementary method. Quantitative PCR were performed on duplicated experiments.

### BioMe WES and EHR data processing and analysis

Genotyping of the whole exome sequencing data from Biome biobank, was described in previous publication ^51,52^. Briefly, Biome is a diverse DNA biobank linked with electric health records from the Mount Sinai Health System (MSHS) based in New York, NY. Patients’ plasma and blood samples was taken for DNA sequencing with their unidentified heath records upon consent. The sample preparation, exome sequencing, quality control and SNP calling were conducted at the Regeneron Genetics Center as described in previous publication ^53^ with GATK best practice. The 4 consecutive SNP within *LILRB3* was not identified while the adjacent SNP rs113314747, was used as an alternative as it is highly linked with those 4 SNPs. The self-reported race information was extracted from the questionnaire of each patient upon enrollment. There are 30,099 patients with WES data and self-reported race including 7,096 African Americans. The disease diagnosis information of each patient was extracted by the International Classification of Disease (ICD)-10 code from EHR records. To assess the association of the SNP association with the severity of kidney diseases, longitudinal eGFR values for the patients with kidney diseases were extracted from the blood test records. Based on the year of birth information of each patient, the eGFR value of each patient at each age was calculated as the mean value of the tests within each year. For patients with more than 3 year-mean-eGFR values less than 15, the first year was marked as the date of eGFR decline and subject to survival analysis.

### Statistical analysis

Cox regression model was used for correlating the expression (alternative allele expression rate) or genotypes of *LILRB3*-4SNPs or *APOL1* G1/G2 to DCGL, in univariate model and adjusted model (adjusted for donor status, HLA mismatch score, induction therapy and recipient genetic race) in GoCAR cohort. The correlation of Banff g score (GoCAR or BioMe), IFTA (CTOT19) or 30% eGFR decline (GoCAR and CTOT19) with *LILRB3*-4SNPs was evaluated by Fisher’s exact test. In BioMe cohort, the clinical diagnosis (defined by ICD10 code) was tested with *LILRB3*-4SNPs and *APOL1* G1/G2 alleles using Logistic regression test. The historical eGFR values were correlated with *LILRB3*-4SNPs or *APOL1* G1/G2 in kidney transplant, ESRD, Non-ESRD and AKI patients using t test.

## Data Availability

The genomic data of pre- and post-transplant specimen in this study were posted to NCBI Gene Expression Omnibus database (GSE252274). The RNAseq data from VericiDx Inc. is available upon request.

## Author’s Contributions

Z.S. performed computational data analysis and was involved in drafting/editing the paper. Z.Y. performed computational data analysis. C.W. led the genomic and functional experiments. W.W. performed genomic and functional experiments. P.C. was involved data interpretation and edited the paper. F.T. and S.W. were involved in pathological and clinical data compiling and interpretation of transplant patients in BioMe cohort. E.A. supervised the serum mass spectrometry profiling. D.S. and Y.L. supervised the analysis of evolutionary selection of LILR locus. S.A. performed immune response assay. T.R. generated *LILRB3*-4SNPs overexpression system. S.L. modeled the interactions of *LILRB3*-SHP2. D.L. guided the ferroptosis experiments. J.F. performed single cell RNAseq. C.X. managed the clinical data. T.L. and H.L. performed mass spectrometry experiments on pretransplant serum. TH.V. performed genetic association analysis.

G.M. performed serum profiling. Q.S. analyzed the evolutionary trait of LILR locus. A.K. and Z.Z. edited the paper. S.F., K.C. were involved data interpretation and edited the paper. J.O. guided the immune functional experiments in THP-1 cells. K.L. edited the paper. S.C. supervised G.M. in serum profiling and edited the paper. J.X. supervised the RNA sequencing. Pa.C. was involved in data generation of CTOT19 and VericiDx cohorts. L.G., R.C. and M.M. were involved data interpretation and edited the paper. G.N. supervised TH.V. in genetic association analysis and edited the paper. C.H. supervised J.F. in single cell RNAseq analysis/data interpretation and edited the paper. M.K. was involved data interpretation and edited the paper. X.J. supervised D.L. in ferroptosis analysis and edited the paper. X.Z. supervised S.L. in structural modeling and edited the paper. WG. Z supervised R. L. in *in vitro LILRB3*-4SNPs overexpression experiments and edited the paper. S.C. supervised S.A in immune response assays and edited the paper. P.H. was involved in study design and data interpretation and edited the paper. W. Z. conceptualized and designed this study and drafted/edited the paper. All authors reviewed and approved the manuscript.

## Acknowledgement

We gratefully acknowledge the administration and IT team of Institute of Personalized Medicine (IPM) at Mount Sinai for facilitating access to genetic and clinical data of BioMe cohort. We also acknowledge the Genomic Resources Core at Well Connell Medical School for generating bulk and single cell RNA sequencing data on GoCAR cohort. We thank VericiDx Inc. for providing the RNAseq dataset of the CTOT19 cohort and RNAseq and demographic data of the VericiDx cohort. We thank Associate IT director, Ricky Kwan, for generating pathological report for transplant patients in BioMe cohort. This work was supported by NIH 5U01AI070107-03 and Biocomputation Fund (02435913) from the Department of Medicine. This work was also supported in part through the computational and data resources and staff expertise provided by Scientific Computing and Data at the Icahn School of Medicine at Mount Sinai and supported by the Clinical and Translational Science Award (CTSA) grant UL1TR004419 from the National Center for Advancing Translational Sciences. Xuewu Zhang is supported in part by grants from the NIH (R35GM130289) and Welch foundation (I-1702). The mass spectrometry data were obtained from an Orbitrap mass spectrometer funded in part by NIH grants NS046593 and 1S10OD025047-01, for the support of proteomics research at Rutgers Newark campus.

## Conflict of Interest Statement

Dr. Zhang reports personal fees from VericiDx and reports the patents (1. Patents US Provisional Patent Application F&R ref 27527-0134P01, Serial No. 61/951,651, filled March 2014. Method for identifying kidney allograft recipients at risk for chronic injury; 2. US Provisional Patent Application: Methods for Diagnosing Risk of Renal Allograft Fibrosis and Rejection (miRNA); 3. US Provisional Patent Application: Method for Diagnosing Subclinical Acute Rejection by RNA sequencing Analysis of a Predictive Gene Set; 4. US Provisional Patent Application: Pretransplant prediction of post-transplant acute rejection.); Dr Menon receives research support from Natera. Dr. Cravedi is a consultant for Chinook therapeutics. Dr. Lorenzo Gallon is the non-executive Director and Chair of the science advisory board for Verici. Other investigators have no financial interest to declare.

