## Supplementary figures and tables for "Genetic polymorphisms of Leukocyte Immunoglobulin-Like Receptor B3 (*LILRB3*) gene in African American kidney transplant recipients are associated with post-transplant graft failure"

### Supplementary methods and files

#### Study Cohorts and related omics datasets:

Our study incorporates three kidney transplant cohorts with each focusing on unique aspects of kidney transplantation, namely GoCAR[1, 2], CTOT19[3], VericiDx, and one EHR-linked biobank cohort (BioMe) [4] as depicted in Figure 1. The GoCAR cohort (genetically-identified 20.6% AA, 63.6% European) was a multi-center kidney transplant study with an aim to uncover genetic and transcriptomic markers associated with diverse transplant outcomes such as acute rejection and renal failure with the period of up to 5-year follow up [1, 2]. The blood samples were collected pretransplant. Kidney biopsies were performed at 3-, 12-, and 24- month surveillance or clinical indications and biopsy core tissues and blood were simultaneously collected. The histological slides were centrally scored by MHG pathology group. All patients underwent genotyping using SNP arrays[5], and RNA sequencing data (150bp pair-ended reads) of pretransplant blood samples from 170 patients [2] (GSE 112927) was used to identify expressional SNPs associated with graft loss. Targeted DNA sequencing of the region covering the identified *LILRB3*-4SNPs was performed on 127 non-European (AA+Hispanic) recipients to validate the *LILRB3*-4SNPs. Mass spectrometry was performed on the serum in pre-transplant blood from 16 AA recipients (8 with and 8 without *LILRB3*-4SNPs) to correlate with RNA sequencing data. Single cell RNA sequencing was performed on the pre-transplant PBMC from 2 patients (1 with and 1 without *LILRB3*-4SNPs) and the post-transplant PBMC at 24 months post-transplant from 2 patients (1 with and 1 without *LILRB3*-4SNPs) to investigate the transcriptomics dysregulation in immune subsets. RNA sequencing data of posttransplant blood samples collected post 6 months biopsies from 20 recipients (10 with and 10 without. *LILRB3*-4SNPs) and biopsy samples collected at 3 months post-transplant from 6 recipients (3 with and 3 without *LILRB3*-

4SNPs) were used to correlate the SNPs with post-transplant gene expression in the posttransplant blood or graft respectively.

The CTOT19 cohort (38.2% AA, 44% White) represents a randomized controlled trial investigating the efficacy of Remicade induction therapy for deceased donor kidney transplant recipients within 2 year followed-up (NCT02495077)[3]. With a cohort of 225 patients, the study spanned 2 years to assess the effectiveness of this therapy. RNA sequencing was executed on pretransplant blood from 128 recipients, utilizing 75bp-long paired-end (75PE) reads. Notably, the recipients were representative of genetically-identified race/ethnic groups: 47 African American, 27 Caucasian, 41 Hispanic, and 13 individuals from other ethnic backgrounds. RNA sequencing data in this cohort was used to validate the variant expression of *LILRB3*-4SNPs in AA recipients. The RNA data in 47 AA recipients was combined with two cohorts to identify the meta-gene signatures associated with the *LILRB3*-4SNPs. The SNPs was also correlated with post-transplant outcomes (eGFR and IFTA within 24 months).

The VericiDx cohort is an ongoing clinical trial (241 patients (32.0% AA) enrolled as of 06/30/2023) with a focus on predicting early acute rejection and renal fibrosis post-transplantation (NCT04727788). Pretransplant RNA sequencing profiles (75PE reads) were generated on pretransplant blood, with a subset of 71 recipients being genetically-identified African American. The RNA data in these 71 AA recipients was combined with two cohorts to identify the meta-gene signatures associated with the *LILRB3*-4SNPs. However, the relatively short follow-up duration for many patients in this cohort limited the potential for robust outcome association validation.

The BioMe cohort, established by the Institute of Personalized Medicine at Mount Sinai, stands as an EHR-linked biobank comprising over 34,000 patients from the Mount Sinai Health system, as of September 30, 2022[4]. Among this cohort, whole exome sequencing (WES) data were obtained from 30,099 patients, including self-reported 7,096 African American, 9,555 Caucasian, 10,467 Hispanic, and 2,981 individuals from other ethnic backgrounds. This data was correlated with EHR information to assess the connection between identified SNPs and laboratory test results, as well as disease diagnoses, within the African American population and compared to that for *APOL1* G1/G2 alleles.

**Detection of expressional SNPs from RNA sequencing data of pretransplant blood**

**Variant genotyping, imputation and filtering**: The quality control, SNP calling and imputation was performed on SNP array data of blood RNA samples from the kidney transplant recipients as described in previous study [5]. The SNP calling from RNAseq data of pretransplant blood samples from the recipients was performed with the GATK best practice [6] under GATK/4.2.0.0. Briefly, sequencing reads were aligned to the human genome hg19 with STAR/2.6.1d [6] using two-pass mode to get better alignments around novel splice junctions. Duplicated reads removing and reads sorting were performed with Picard/1.93. SplitNCigarReads from GATK was used to splits reads with N in the cigar into multiple supplementary alignments and hard clips mismatching overhangs for the specific alignment for RNAseq data. Base quality was recalculated with BaseRecalibrator and variants were called with HaplotypeCaller. Variants from different samples were merged together with GenotypeGVCFs and variants with FS > 30.0 and QD < 2.0 were considered as low quality and filtered. Finally, we combined the genotypes from SNP array and RNAseq, following the order of the confidence level as Array > imputation > RNAseq.

**Quantification of allele specific expression**: The SNP genotypes from SNP arrays and RNAseq were then combined and annotated with AnnoVar version "2018Apr16". The exonic

SNPs were subsequently extracted for further analysis. To quantify the RNA sequencing reads that are mapped to these exonic SNPs, the nucleotides of those exonic SNP locus on human genome (hg19) were masked as "N" to avoid mapping bias towards reference allele. RNA sequencing reads were then aligned to the masked human genome with STAR 2.6.1d and duplicate reads were identified by Picard 1.93. The reads mapped to reference and alternative alleles of each SNP were counted with ASEReadCounter. The alternative allele expression fraction (AEF) of each SNP was calculated as read count for alternative allele / (the total read count for reference and alternative allele). The reads covering target SNP site was extracted with samtools 1.9 and the coverage was visualized with IGV 2.8.2.

The same procedure was also applied to the RNA sequencing dataset from CTOT19 and VericiDx cohorts to detect the allelic specific expression of LILR3-4SNPs identified in GoCAR cohort.

**Targeted DNA sequencing of the LILR3-4SNPs locus:**

The DNA samples from 124 non-Caucasian (AA+Hispanic) recipients in GoCAR cohort were retrieved from GoCAR specimen biobank and were subjected to targeted DNA sequencing by following Illumina 16S sequencing protocol. The DNA region covering the *LILRB3*-4SNPs was amplified using the primers (5′-TCGTCGGCAGCGTCAGATGTGTATAAGAGACAGTTCTGCTGAGTGTGGGGTCT-3′ and 5′- GTCTCGTGGGCTCGGAGATGTGTATAAGAGACAGGCCTCCCAGGATGTGACCTA-3′) with addition of the Illumina adaptor overhangs. The PCR amplification was performed in a total 20 μl reaction volume for each DNA sample. After purification of amplicons using AMPure XP beads (Beckman) and confirmation of successful amplification by high sensitivity DNA assay for Bioanalyzer, the amplification product was proceeded to attach dual indices and Illumina sequencing adapters using the Nextera XT Index Kit (Illumina) to PCR product for each DNA sample. Library quantification was performed using the Qubit dsDNA HS Kit (Thermo Fisher Scientific), and the size distribution was assessed using a high-sensitivity DNA assay for Bioanalyzer and qualified libraries were then pooled for sequencing. Sequencing of pooled libraries was performed on the NovaSeq 6000 platform (Illumina) at paired-end 150bp with 20M reads.

To determine genotypes of the targeted region around the LILR3-4SNPs locus, the reference sequences of 200pb upstream and 200bp downstream rs549267286 was extracted from human genome reference (hg19) to build the targeted reference genome using bwa/0.7.15. The DNA sequences were aligned to the targeted reference with bwa mem with default parameters. SNPs within this region were called by bcftools/1.9 with mpileup by default parameters for each sample and finally merged by bcftools merge.

**Race disparity and natural section of *LILRB3*-4SNPs**

Race disparity: The frequency of *LILRB3*-4SNPs in the population with different races was independently assessed in multiple cohorts. (RNA sequencing dataset of CTOT19, VericiDx and Sclerosis, targeted sequencing data of *LILRB3*-4SNPs locus from non-European population in GoCAR cohort and whole exome sequencing data from BioMe cohort)

Natural selection: We used the 1000 Genomes Selection Browser (https://www.ncbi.nlm.nih.gov/pmc/articles/PMC3965045/) to examine ranked scores for SweepFinder's CLR test statistic, which detects skews in allele frequency at varying distances from a selective sweep, Fay and Wu's H, which highlights regions with an excess of high-frequency derived alleles that are expected after a sweep, pairwise FST, which measures genetic differentiation between two population samples and which will be elevated when either population has experienced a selective sweep not shared with the other, and XP-CLR, which extends the concept of the SweepFinder CLR to pairwise population comparisons in order to detect population-specific sweeps. For each of these statistics, elevated rank scores are potentially indicative of selective sweeps.

**Serum mass spectrometry of pretransplant serum**

The relative abundances of serum proteins in pretransplant blood of selected 16 AA recipien were quantified using mass spectrometry (MS)-based proteomics. Preprocessing of 20 µl of each serum sample was carried out using the Enrich-iST preparation kit, following the manufacturer's protocol with a slight modification of conducting an overnight digestion at 37ºC. Specifically, 20 µl of serum from each sample was mixed with magnetic beads (En-Beads) for 30 minutes on a thermoshaker at 30°C with 1200 rpm. The proteins binding on beads were then reduced, alkylated, followed by on-beads trypsin digestion. The resulting peptides were desalted and analyzed using an Orbitrap Fusion Lumos Tribrid mass spectrometer (Thermo Scientific) as previously described [7]. Briefly, peptides were separated for two hours on C18 columns using a binary gradient of 2% acetonitrile in 0.1% formic acid and 85% acetonitrile in 0.1% formic acid. Peptides were then introduced into the mass spectrometer via nanospray Flex ion source. The spectra were acquired in a positive mode at 2 kV and 275ºC, scanning within a range of m/z 375 to 1500 with 120,000 FWHM on an orbitrap. Peptides with charge states 2-7 were selected for further MS/MS analysis. The acquired spectra were then searched against the Uniprot validated human proteome (downloaded on September 11, 2023) using Sequest search engine through Proteome Discoverer (v. 2.4). Mass tolerance was 10 ppm for MS and 0.6 Da for MS2. Methionine oxidation and acetylation of N-termini were set as variable modifications, and cysteine carbamidomethylation was set as a fixed modification. Maximum allowed false discovery rate was 1%.

The expression of each protein between samples with and without risk allele was tested using Student's T test. Proteins with p value < 0.05 was identified as differentially expressed (DEP). WebCSEA [8] was used to predict the enriched cell type of up regulated (log2(fold change) > 0) and down regulated (log2(fold change) < 0) DEP separately.

**Single cell RNA sequencing analysis on pre- and post-transplant PBMC**

The single-cell RNA sequencing library was prepared for 2 pre- and 2 post-transplant PBMC samples from the GoCAR cohort, and each condition has one sample with and one sample without *LILRB3*-4SNP risk allele, whose genotype was identified in SNP array or RNAseq data as described above.

The pre- and post-transplant PBMC sequencing data were analyzed separately as they were sequenced in different batch. The raw sequencing data was aligned to human genome hg38 and quantify gene expression with cellranger (7.1.0) count with default parameters. The gene-cell count matrix was submitted to Seurat 4.1.1 [9] for QC, normalization and clustering. Genes expressed in less than 3 cells were removed and cells expressing less than 200 genes and more than 7000 genes were removed as low quality and potential doublets. Cells with more than 30% RNA expressed from mitochondrial genes were also removed as low quality. The gene expression of each cell was normalized with its total expression and multiplied with scale factor 10000 and then log-transformed. Harmony was used for batch correction between individuals with function RunHarmony [10]. Then unsupervised clustering was performed with FindNeighbours with the first 30 "harmony" dimensions and FindClusters with resolution of 0.8 with the default method Louvain algorithm. FindMarkers was used to identify the markers of each cluster by comparing each gene with other clusters with Wilcoxon Rank Sum test. Each cluster was annotated with classic markers for each cell type in PBMC as published previously [11]. The differentially expressed genes (DEG) between risk and non-risk sample of each cell type was identified with FindMarkers with Wilcoxon Rank Sum test. Genes with p value < 0.001, log2(fold change) <= 0.25 or >= 0.25 and expressed in at least 10 percent of cells in each condition were identified as differentially expressed. Function enrichment analysis of DEG was performed with Enrichr [12] with upregulated (log2(fold change) > 0) and downregulated (log2(fold change) < 0) genes separately.

**The primer sets of inflammation and ferroptosis related genes for qPCR assay**

The following primer sets used were synthesized by Sigma-Aldrich:
human GAPDH (Forward, 5′-AATTGAGCCCGCAGCCTCCC-3'; Reverse, 5′-CCAGGCGCCCAATACGACCA-3'),
TNF-α (Forward, 5′-ACGCTCTTCTGCCTGCT-3'; Reverse, 5′-GCTTGAGGGTTTGCTACAA-3'),
TNFAIP3F (Forward, 5′- GAAGAAACTCAACTGGTGTCG -3'; Reverse, 5′-CCAAGTCTGTGTCCTGAACG-3'),
IL1-β (Forward, 5′- TGGCTTATTACAGTGGCAATG-3'; Reverse, 5′-TGGTGGTCGGAGATTCGT-3'),
CCL2 (Forward, 5′- GAATCACCAGCAGCAAGTG-3'; Reverse, 5′-CTTCGGAGTTTGGGTTTG-3'),
BCL3 (Forward, 5′- CTTTCTGCTGACATCGCC-3'; Reverse, 5′-GTCTGCCGTAGGTTGTTGTA-3'),
NFkB1AF (Forward, 5′- GAAGGCTACCAACTACAATGG-3'; Reverse, 5′-TTCAACAGGAGTGACACCAG-3'),
GPX4 (Forward, 5′- GCCTTCCCGTGTAACCAGT-3'; Reverse, 5′-GCGAACTCTTTGATCTCTTCGT-3'),
FTL (Forward, 5′- CAGCCTGGTCAATTTGTACCT-3'; Reverse, 5′-GCCAATTCGCGGAAGAAGTG-3'),
NCOA4 (Forward, 5′- GAGGTGTAGTGATGCACGGAG-3'; Reverse, 5′-GACGGCTTATGCAACTGTGAA-3'),
FTH1 (Forward, 5′- CCCCCATTTGTGTGACTTCAT-3'; Reverse, 5′-GCCCGAGGCTTAGCTTTCATT-3'),
TFRC (Forward, 5′- AAAATCCGGTGTAGGCACAG-3'; Reverse, 5′-GCACTCCAACTGGCAAAGAT-3'),
ACSL3 (Forward, 5′- GCCGAGTGGATGATAGCTGC-3'; Reverse, 5′-ATGGCTGGACCTCCTAGAGTG-3').

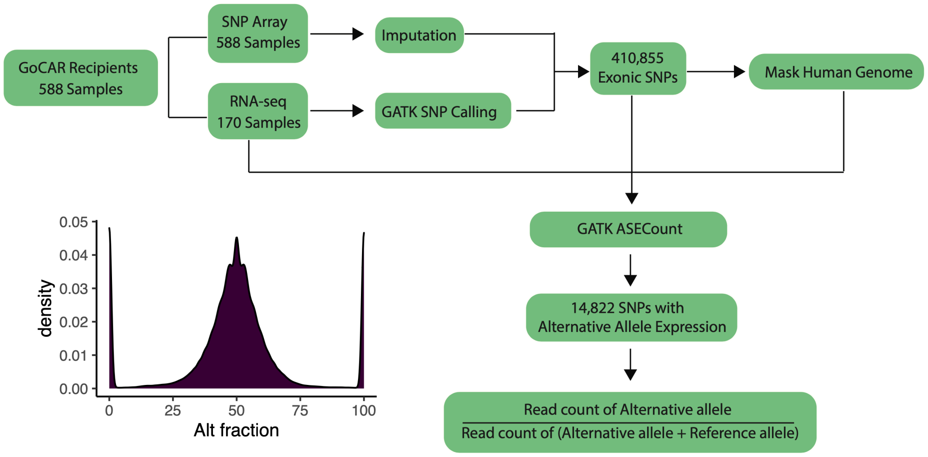


**Supplementary Figure 1.** Workflow of the quantification of allele specific expression in GoCAR cohort.

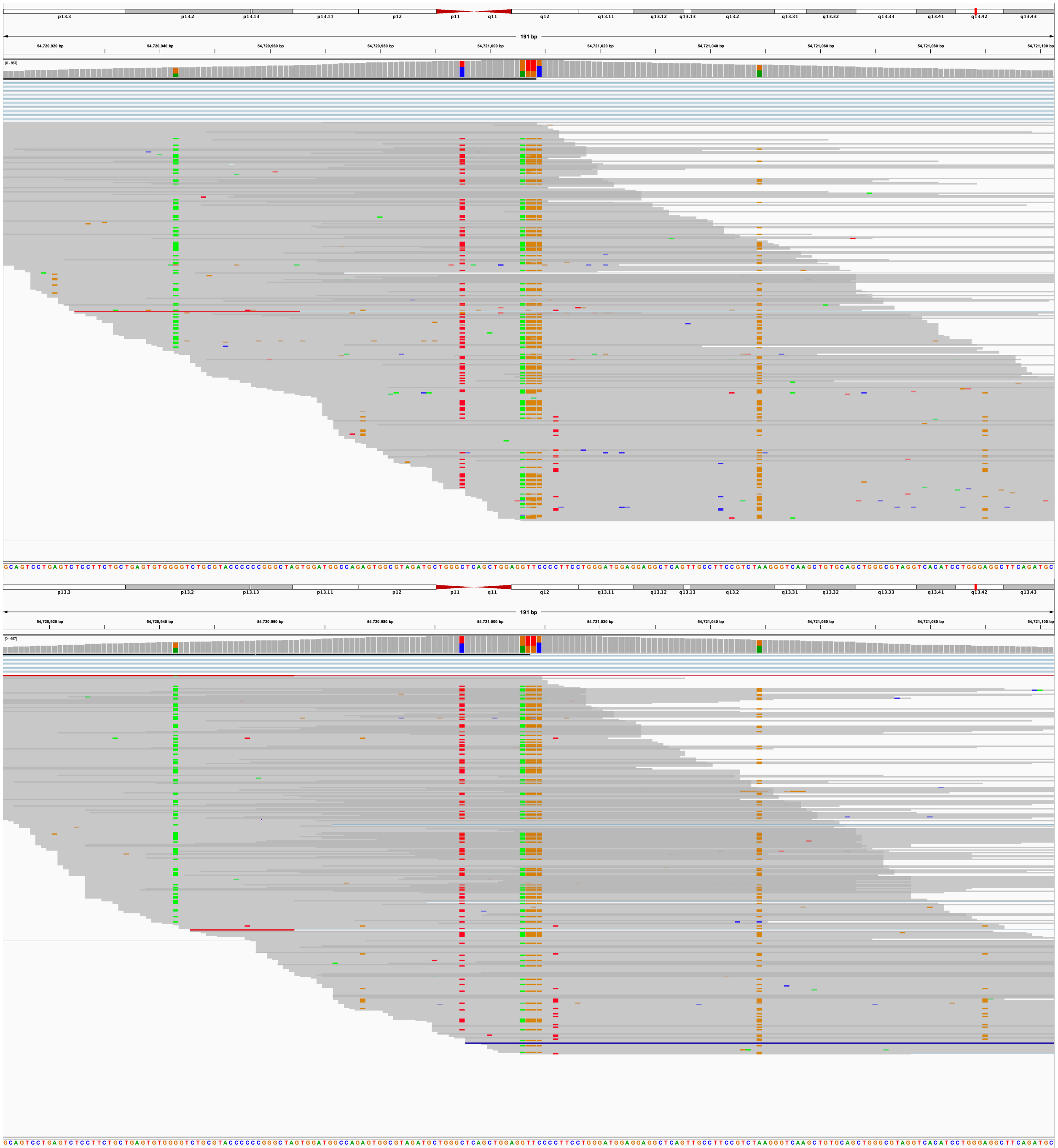

**Supplementary Figure 2** IGV plot of *LILRB3*-4SNPs in 2 different patients, P15313 (upper) and P36683 (Lower).

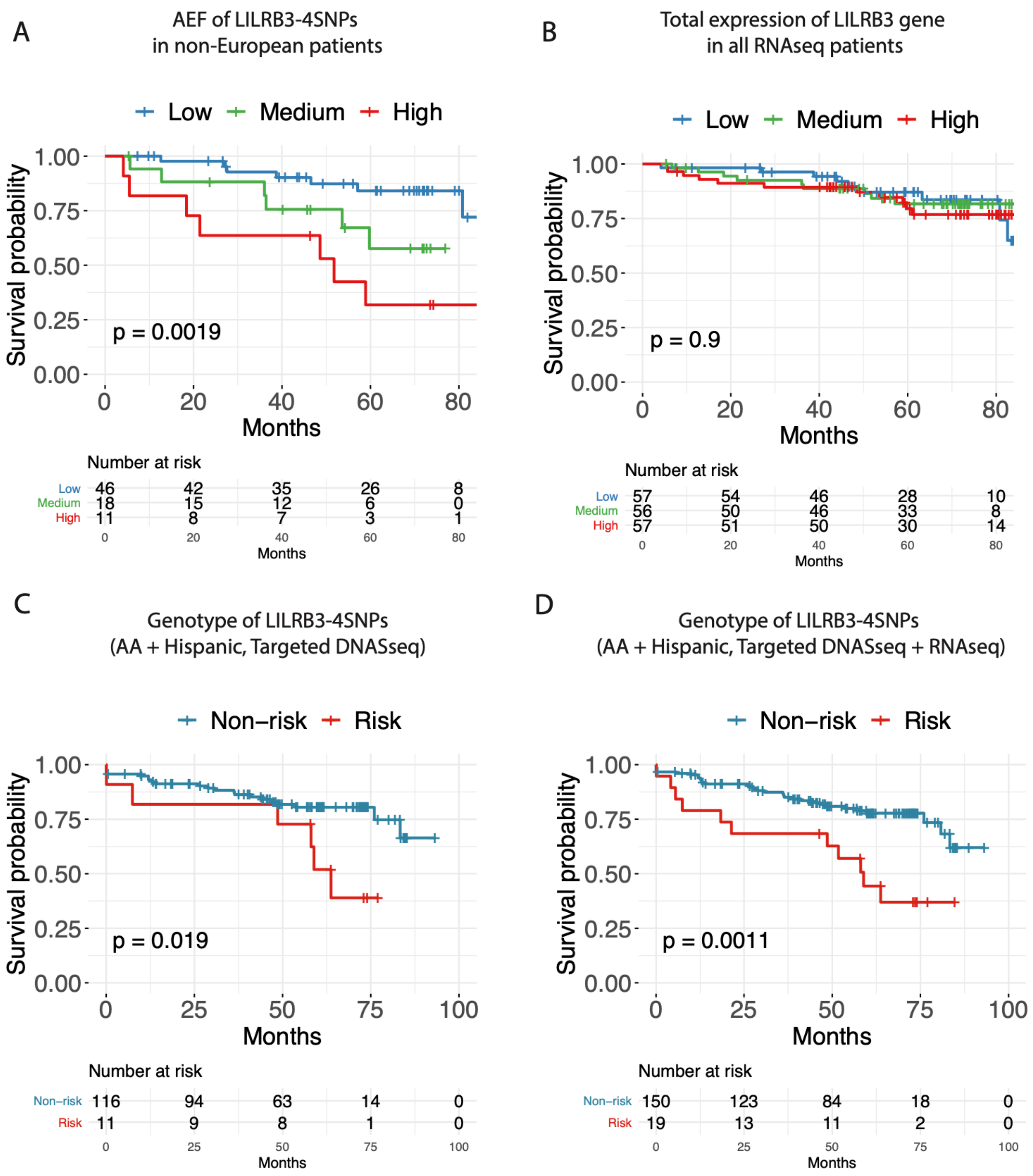


**Supplementary Figure 3.** Association of the AEF of *LILRB3*-4SNP in non-European (AA + Hispanic, n = 75) (**A**), total expression of *LILRB3* in all RNAseq cohort (n = 170) (**B**), genotype of *LILRB3*-4SNPs in non-European targeted sequencing cohort (AA + Hispanic, n = 127) (**C**) and targeted sequencing + RNAseq cohort (AA + Hispanic, n = 169) (**D**) with DCGL in the GoCAR cohort.

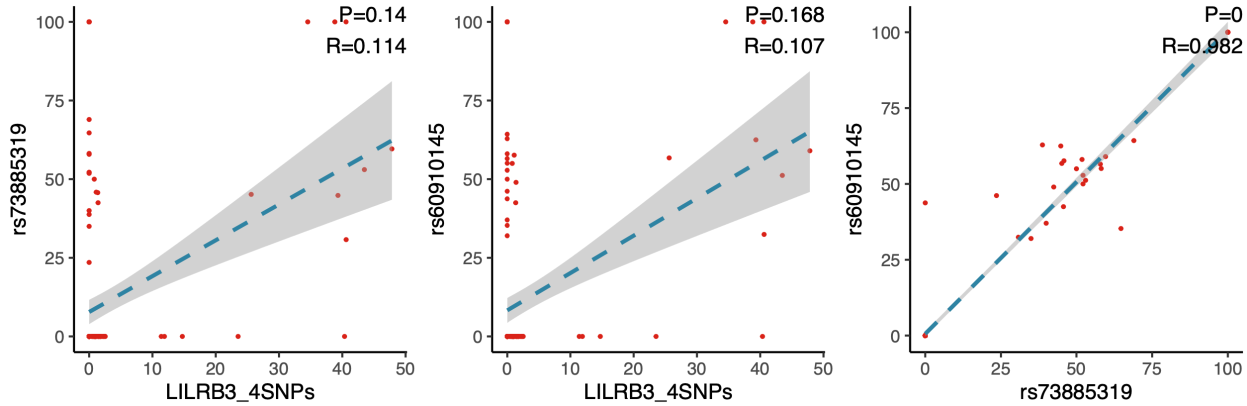


**Supplementary Figure 4.** Correlation (Spearman correlation) between *LILRB3*_4SNPs the AFE of *APOL1* G1 allele (rs73885319 (**left**) and rs60910145 (**middle**)), and two *APOL1* G1 alleles (**right**).

| Population | Group | Sample Size | Ref Allele | Alt Allele |
|---|---|---|---|---|
| **Total** | Global | 153692 | T=0.998543 | G=0.001457 |
| European | Sub | 134352 | T=0.999985 | G=0.000015 |
| African | Sub | 4122 | T=0.9495 | G=0.0505 |
| African Others | Sub | 162 | T=0.920 | G=0.080 |
| African American | Sub | 3960 | T=0.9508 | G=0.0492 |
| Asian | Sub | 6228 | T=1.0000 | G=0.0000 |
| East Asian | Sub | 4438 | T=1.0000 | G=0.0000 |
| Other Asian | Sub | 1790 | T=1.0000 | G=0.0000 |
| Latin American 1 | Sub | 442 | T=0.991 | G=0.009 |
| Latin American 2 | Sub | 950 | T=1.000 | G=0.000 |
| South Asian | Sub | 270 | T=0.996 | G=0.004 |

**Supplementary Figure 5.** Minor allele frequency of rs549267286 (one of *LILRB3*-4SNPs) among different continent population in dbSNP.

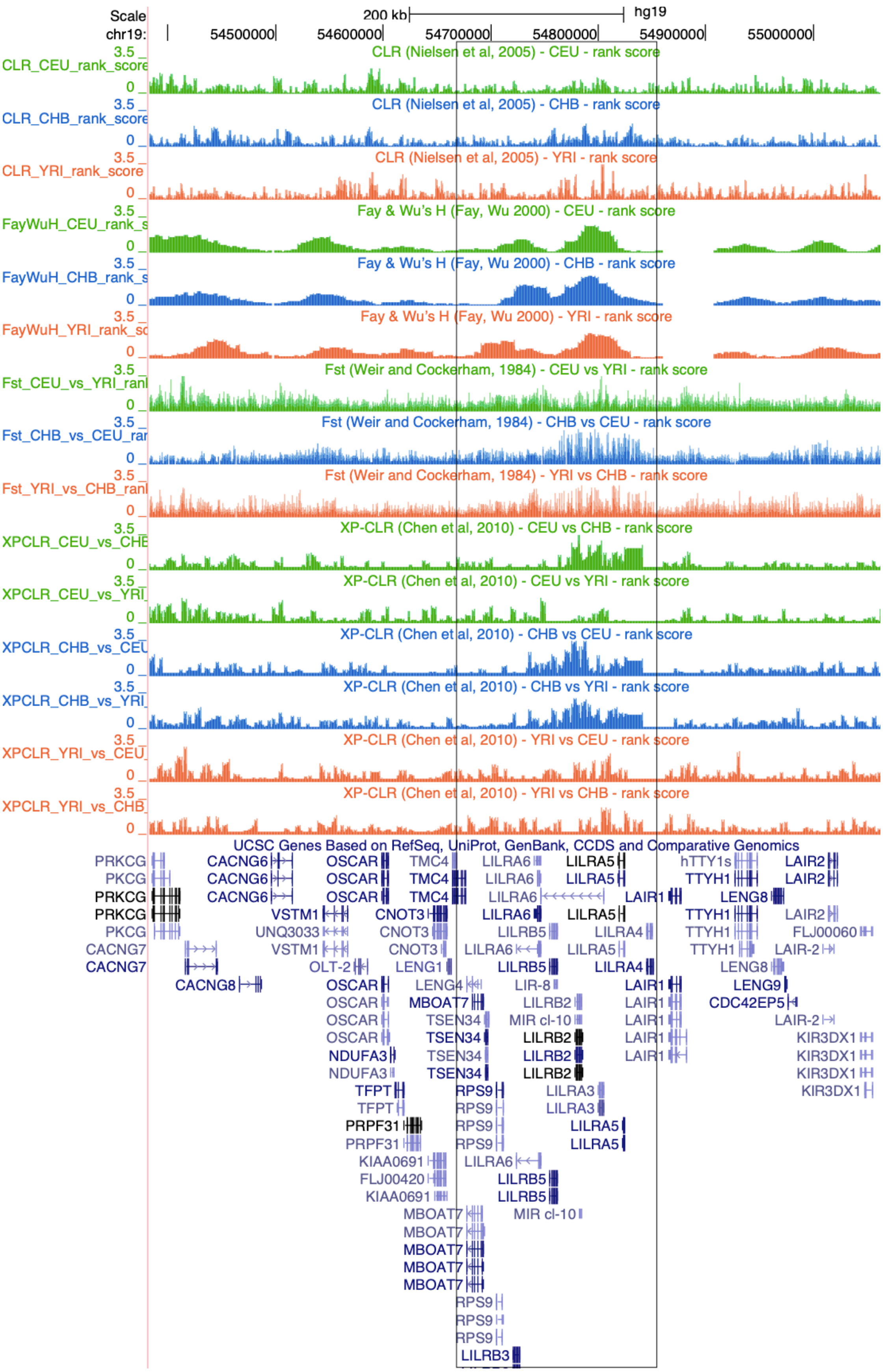
Scale
200 kb
hg19
chr19:
54500000
54600000
54700000
54800000
54900000
55000000
CLR (Nielsen et al, 2005) - CEU - rank score
CLR (Nielsen et al, 2005) - CHB - rank score
CLR (Nielsen et al, 2005) - YRI - rank score
Fay & Wu's H (Fay, Wu 2000) - CEU - rank score
Fay & Wu's H (Fay, Wu 2000) - CHB - rank score
Fay & Wu's H (Fay, Wu 2000) - YRI - rank score
Fst (Weir and Cockerham, 1984) - CEU vs YRI - rank score
Fst (Weir and Cockerham, 1984) - CHB vs CEU - rank score
Fst (Weir and Cockerham, 1984) - YRI vs CHB - rank score
XP-CLR (Chen et al, 2010) - CEU vs CHB - rank score
XP-CLR (Chen et al, 2010) - CEU vs YRI - rank score
XP-CLR (Chen et al, 2010) - CHB vs CEU - rank score
XP-CLR (Chen et al, 2010) - CHB vs YRI - rank score
XP-CLR (Chen et al, 2010) - YRI vs CEU - rank score
XP-CLR (Chen et al, 2010) - YRI vs CHB - rank score
UCSC Genes Based on RefSeq, UniProt, GenBank, CCDS and Comparative Genomics

**Supplementary Figure 6.** Genome browser shot showing signatures of selection in the LILR region ( in Box) in the hg19 coordinate space. This image was created using the 1000 Genomes Selection Browser (https://www.ncbi.nlm.nih.gov/pmc/articles/PMC3965045/). For each statistic, we show the Selection Browser's rank scores, which are calculated as $-\log_{10}(p)$ where p is the percentile ranking of a given statistic in the focal region (taken after ordering values from most- to least-suggestive of positive selection). Thus, values above 2 represent the top 1% of all scores across the genome, and values about 3 represent the top 0.1%. We show ranked scores for SweepFinder's CLR test statistic, Fay and Wu's H, pairwise $F_{ST}$, and XP-CLR (all described in the Methods). For each of these statistics, elevated rank scores are potentially indicative of selective sweeps.

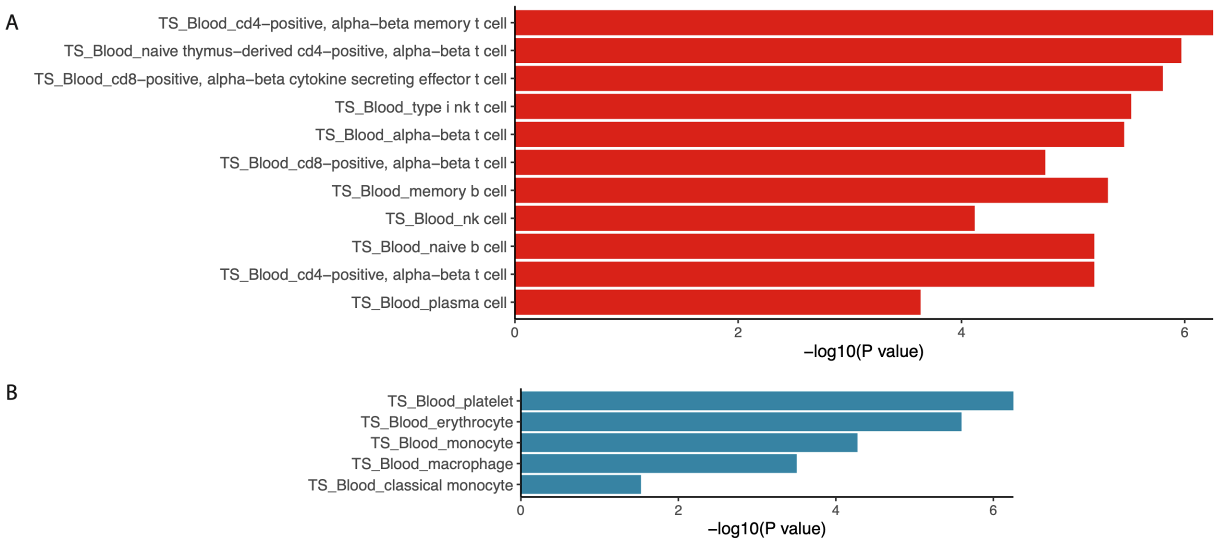


**Supplementary Figure 7**. Enriched blood cell types of up- (n = 892) and down- (n = 663) regulated meta genes among 3 transplant cohort ($p < 0.05$), from webCSEA [35610053].

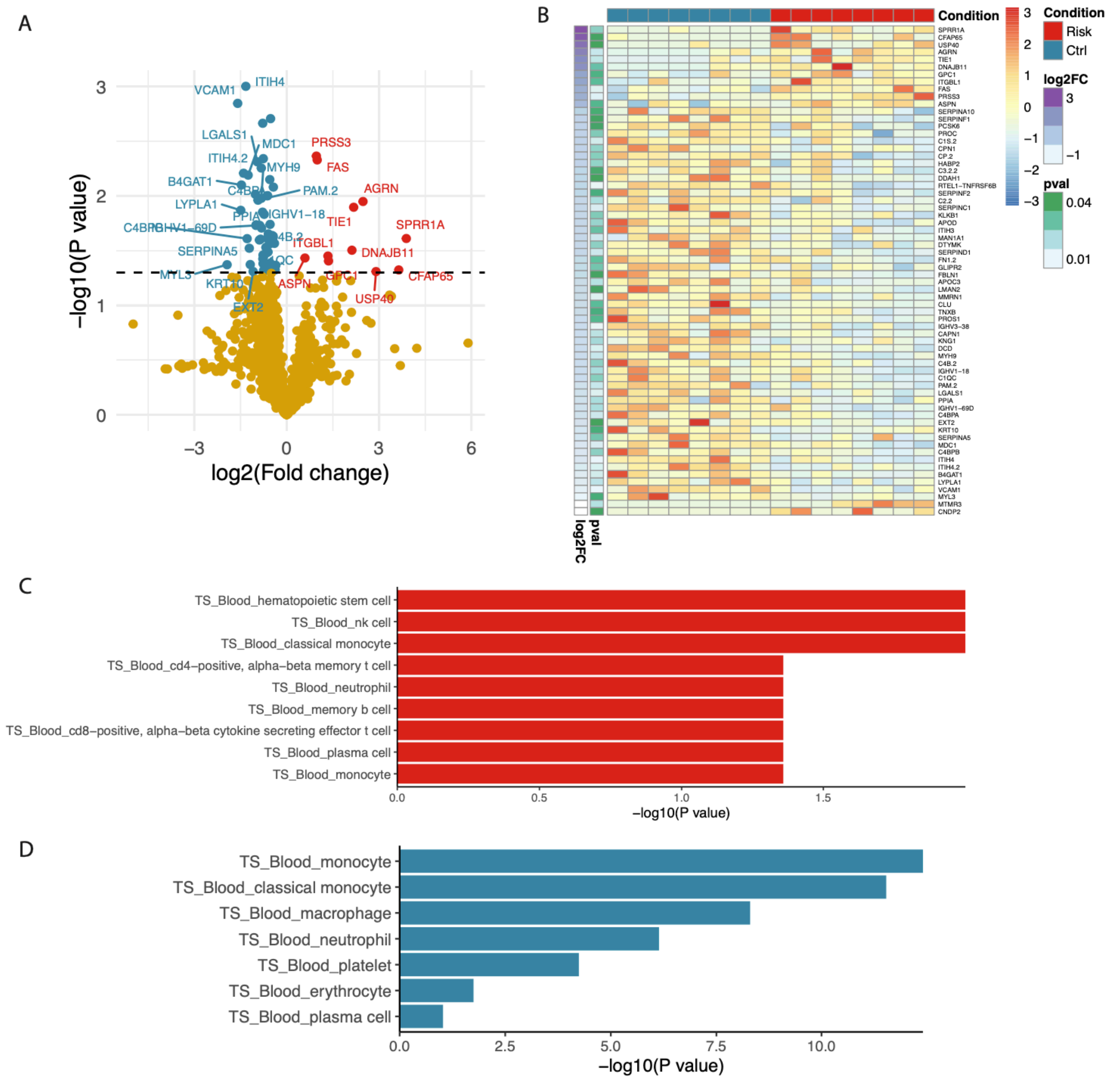


**Supplementary Figure 8.** Differentially expressed serum proteins between *LILRB3*-4SNPs positive (n = 8) and negative patients (n = 8) measured by Mass spectrometry in pre-transplant patients in GoCAR. The significantly dysregulated DEPs were showed by volcano plot (A) or heatmap (B). The cell specificity of up- and down-regulated DEPs was illustrated in (C) and (D), respectively.

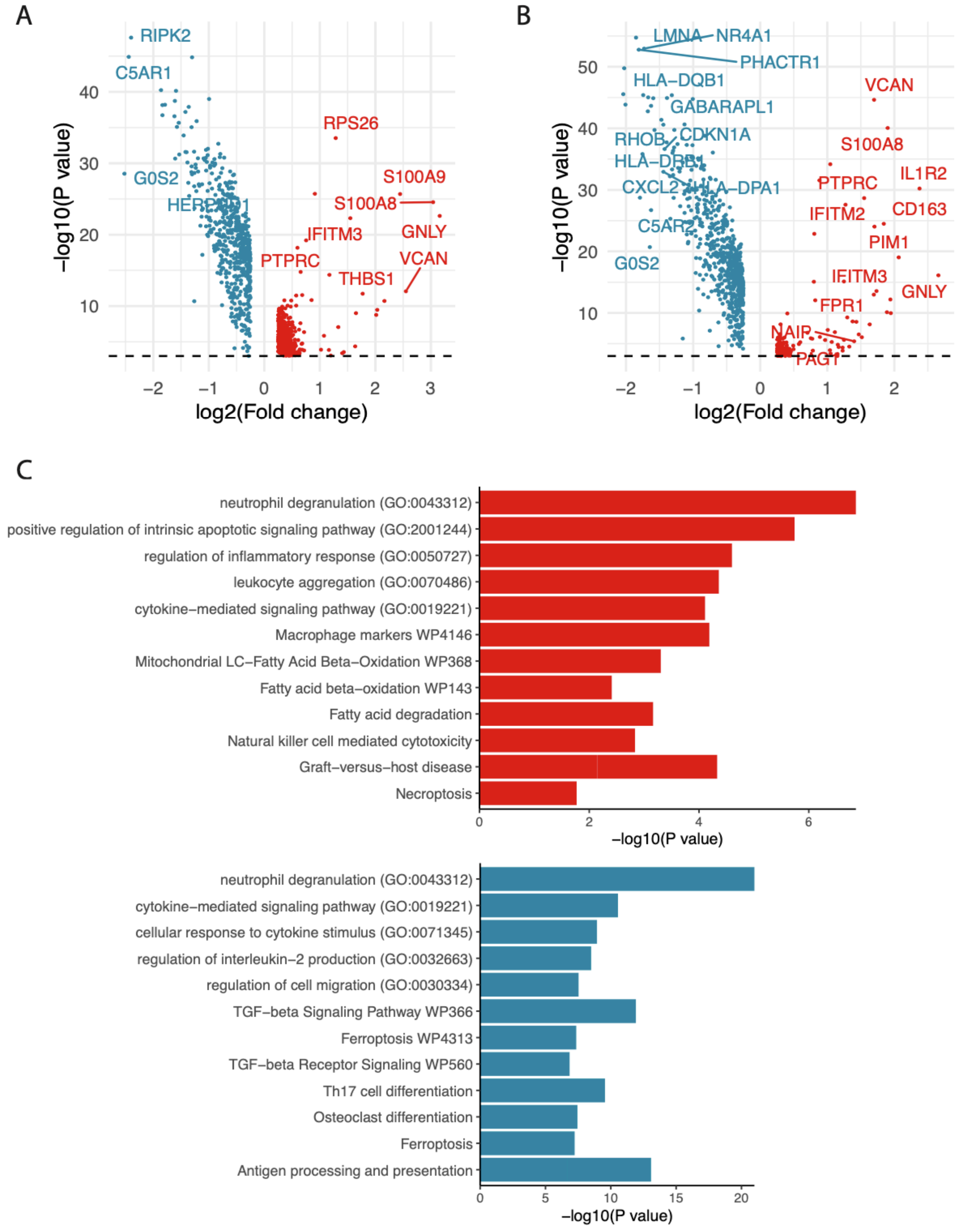


**Supplementary Figure 9.** DEG and function enrichment of CD16+ (A) and CD14+ monocytes (B-C) between two pretransplant recipients with and without *LILRB3*-4SNPs.

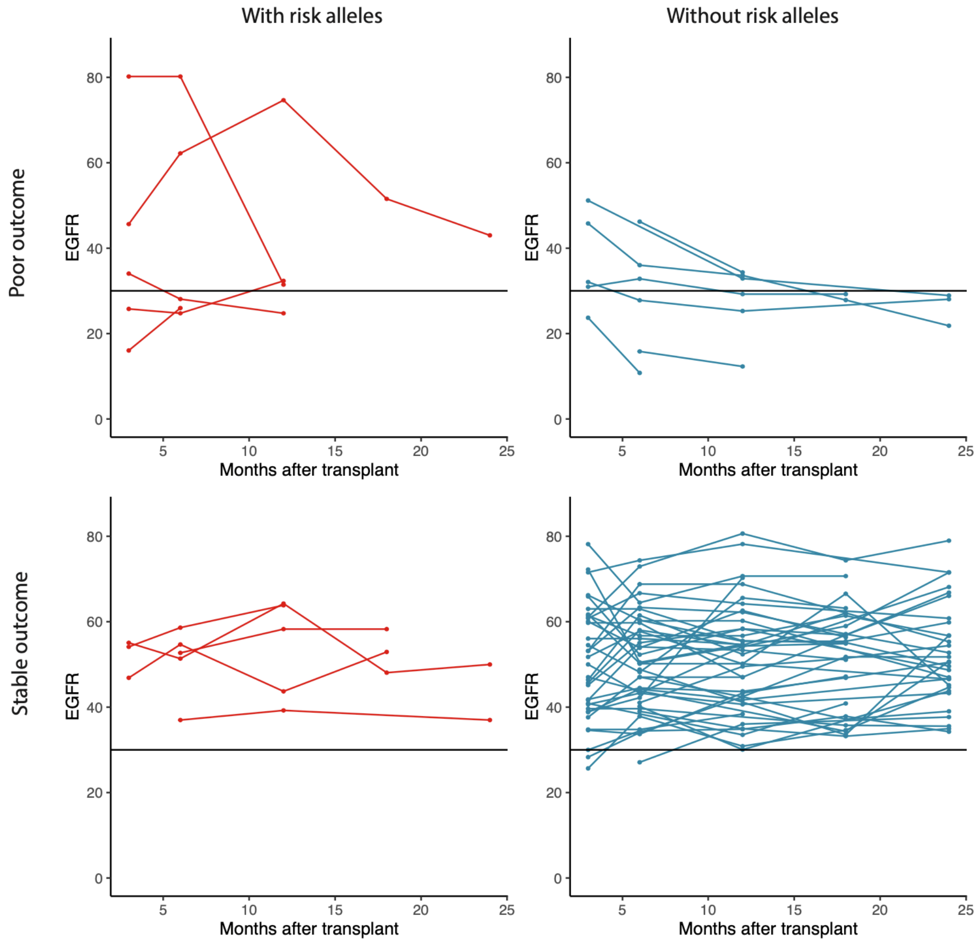


| | Risk | Non-risk |
|---|---|---|
| **Poor** | **5** | **7** |
| **Stable** | **5** | **46** |

*P=0.017 (Fisher's exact test)

**Supplementary Figure 10.** Trend of eGFR (declined in upper panel, and stable in lower panel) in the AA patients with (left) and without (right) *LILRB3*-4SNP (P = 0.01) in the GoCAR cohort.

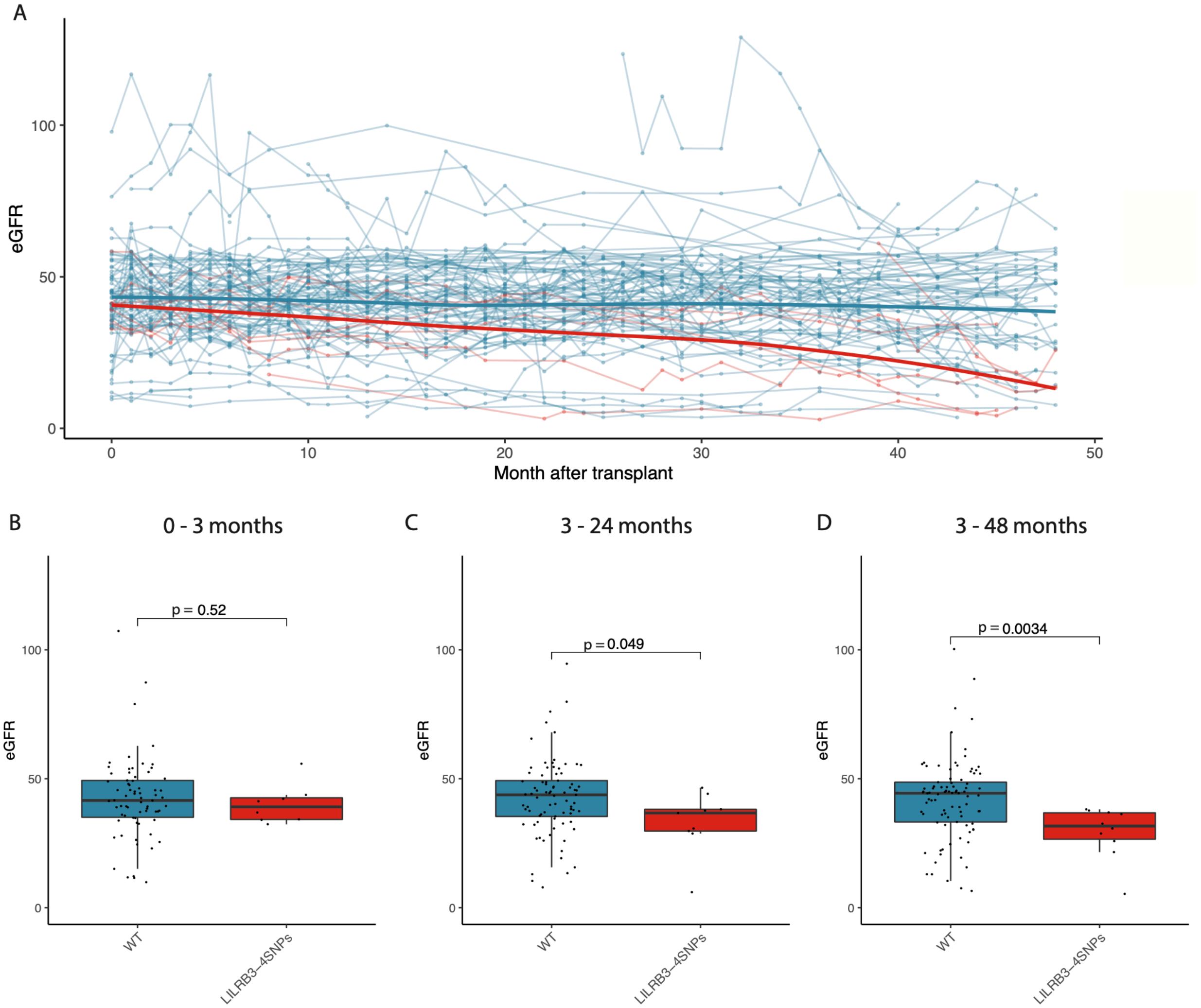


**Supplementary Figure S11. Post-transplant longitudinal eGFR values of AA kidney transplant patients (ICD Z94.0) in the BioMe cohort. A**) longitudinal eGFR values of AA kidney transplant patient with (red curve) and without (blue curve) *LILRB3*-4SNPs within 48 months after transplantation. Bold curves indicate the regression lines for two groups. Comparison (Students' T test) of average eGFR between patients with (n = 9) or without (n = 90) *LILRB3*-4SNPs within 3 months (**B**), 3-24 months (**C**), and 3-48 months (**D**) after transplantation.

**Table S1. Demographic information of transplantation recipients in GoCAR, CTOT19 and VericiDx.**

| | GoCAR (n = 264) | CTOT19 (n = 128) | VericiDx (n = 77) |
|---|---|---|---|
| **Recipient information** | | | |
| Recipient gender (male) | 189/264 (71.6%) | 75/128 (58.6%) | 44/77 (57.1%) |
| Recipient age | 50.84 ± 13.74 | 53.76 ± 11.15 | 52.47 ± 12.91 |
| Recipient race (Genetic race) | | | |
| African American | 83/264 (31.4%) | 47/128 (36.7%) | 77/77 (100%) |
| European | 82/264 (31.1%) | 26/128 (20.3%) | 0/77 (0) |
| Hispanic | 86/264 (32.6%) | 41/128 (32.0%) | 0/77 (0) |
| Unknown | 13/264 (4.9%) | 14/128 (10.9%) | 0/77 (0) |
| **Transplant outcome** | | | |
| Death censored graft loss[a] | 52/264 (19.7%) | 5/128 (3.9) | - |
| Graft loss time (month) | 54.9 ± 23.8 | 17.0 ± 15.8 | - |
| **Underlying disease** | | | |
| Diabetes | 36/221 (16.3%) | 31/128 (24.2%) | 2/77 (2.6%) |
| Diabetes Hypertension | 53/221 (24%) | - | 29/77 (37.7%) |
| Hypertension | 65/221 (29.4%) | 30/128 (23.4%) | 38/77 (49.4%) |
| Glomerular disease | - | 21/128 (16.4%) | 3/77 (3.9%) |
| Other | 67/221 (30.3%) | 46/128 (35.9%) | 5/77 (6.5%) |
| **Induction therapy** | | | |
| Induction | 213/264 (79.7%) | 128/128 (100%) | - |
| No induction | 51/264 (19.3%) | 0/0 (0%) | - |
| **Donor information** | | | |
| Donor age | 42.43 ± 15.15 | 40.54 ± 13.08 | - |

| | | | |
|---|---|---|---|
| Donor gender | 147/264 (55.7%) | 69/128 (53.9%) | - |
| Donor status | | | |
| Living donor | 102/264 (38.6%) | 0/0 (0%) | - |
| Diseased donor | 162/264 (61.4%) | 128/128 (100%) | - |
| Donor race | | | |
| African American | 27/219 (12.3%) | 18/128 (14.1%) | - |
| Caucasian | 134/219 (61.2%) | 86/128 (67.2%) | - |
| Hispanic | 58/219 (26.5%) | 24/128 (18.6%) | - |
| Other | | | |
| HLA mismatch score | 0: 24/264 (9.1%)<br>1: 31/264 (11.7%)<br>2: 97/264 (36.7%)<br>3: 112/264 (42.4%) | Eplet risk low: 34/128(26.6%)<br>Eplet risk medium: 49/128(38.3%)<br>Eplet risk high: 44/128(34.4%) | |

[a]: The follow up time for CTOT19 is 24 months.

**Table S5. Demographic information between African American patients with or without rs549267826 risk allele.**

| | All (n = 83) | Non-risk (n = 67) | Risk (n = 16) | P value |
|---|---|---|---|---|
| **Basic information** | | | | |
| Recipient gender | 56 (67.5) | 43 (66.2) | 11 (68.8) | 1 |
| Recipient age | 52.88 ± 11.71 | 53.62 ± 11.68 | 51.06 ± 11.86 | 0.45 |
| Recipient BMI | 36.19 ± 49.66 | 36.08 ± 51.83 | 37.4 ± 44.84 | 0.92 |
| Transplant Outcome | | | | |
| Death censored graft loss | 29 (34.9) | 18 (27.7) | 11 (68.8) | **0** |
| Graft loss time (month) | 49.93 ± 24.82 | 51.44 ± 24.07 | 41.03 ± 26.83 | 0.17 |
| Underlying disease | | | | **0.09** |
| Diabetes | 10/77 (13) | 10/62 (16.1) | 0/15 (0) | |
| Diabetes Hypertension | 25/77 (32.5) | 22/62 (35.5) | 3/15 (20) | |
| Hypertension | 35/77 (45.5) | 26/62 (41.9) | 9/15 (60) | |
| Other | 7/77 (9.1) | 4/62 (6.5) | 3/15 (20) | |
| Induction therapy | | | | |
| LD | 70 (84.3) | 57 (85.1) | 13 (81.2) | 0.7 |
| LND | 13 (15.7) | 10 (14.9) | 3 (18.8) | 0.7 |
| **Donor information** | | | | |
| Donor_Gender | 53 (63.9) | 41 (61.2) | 12 (75) | 0.56 |
| Donor_Age | 42.19+-16.05 | 41.83+-16.63 | 43.69 ± 13.72 | 0.74 |
| Donor status | | | | 1 |
| LD | 17 (20.5) | 14 (20.9) | 3 (18.8) | |
| DD | 66 (79.5) | 53 (79.1) | 13 (81.2) | |
| Donor race | | | | 0.7 |
| African American | 20/64 (31.2) | 16/51 (31.4) | 4/13 (30.8) | |
| Caucasian | 29/64 (45.3) | 22/51 (43.1) | 7/13 (53.8) | |
| Hispanic | 15/64 (23.4) | 13/51 (25.5) | 2/13 (15.4) | |
| HLA mismatch score | | | | 0.85 |
| 0 | 3 (3.6) | 3 (4.5) | 0 (0) | |
| 1 | 6 (7.2) | 5 (7.5) | 1 (6.2) | |
| 2 | 27 (32.5) | 22 (32.8) | 5 (31.2) | |
| 3 | 47 (56.6) | 37 (55.2) | 10 (62.5) | |

#### Table S6. Linkage of *APOL1* G1/G2 and *LILRB3*-4SNPs genotypes in Biome Biobank.

| | | *APOL1* genotype[a] | | | | | |
|---|---|---|---|---|---|---|---|
| | | WT/WT | WT/G1 | WT/G2 | G1/G1 | G1/G2 | G2/G2 |
| *LILRB3*-4SNPs genotype | 0/0 | 2781 | 1713 | 1048 | 363 | 371 | 136 |
| | 0/1 | 233 | 159 | 92 | 32 | 46 | 6 |
| | 1/1 | 15 | 7 | 8 | 2 | 0 | 1 |

[a]: $R^2$ = 6.7e-05, D' = 0.02 and p value = 0.333 between *APOL1* G1 allele and *LILRB3*-4SNPs.
$R^2$ = 6.69e-06, D' = 0.005 and p value = 0.759 between *APOL1* G2 allele and *LILRB3*-4SNPs.

**Table S7. Cox model of *LILRB3*-4SNPs genotype associating with DCGL within AA patients adjusted by *APOL1* genotype.**

| | 95% CI | Hazard ratio | P value |
|---|---|---|---|
| *LILRB3*-4SNPs | (0.29, 2.25) | 3.57 | **0.011** |
| *APOL1* single allele | (-0.70, 2.52) | 2.47 | 0.270 |
| *APOL1* double alleles | (-0.73, 2.35) | 2.24 | 0.304 |

**Table S8. Cox model and logistic model of *LILRB3*-4SNPs genotype associating with DCGL within AA patients adjusted by induction therapy, donor status and HLA mismatch.**

| | %95 confident interval | Hazard (Odds) ratio | P value |
|---|---|---|---|
| **Cox model** | | | |
| *LILRB3*-4SNPs | (1.59 8.04) | 3.58 | 0.002 |
| Donor status | (1.26, 25.29) | 5.65 | 0.024 |
| HLA mismatch | (0.45, 1.22) | 0.74 | 0.242 |
| **Logistic Model** | | | |
| *LILRB3*-4SNPs | (2.15, 30.51) | 7.38 | 0.003 |
| Donor status | (1.58, 67.55) | 7.94 | 0.026 |
| HLA mismatch | (0.36, 1.61) | 0.77 | 0.472 |

**Table S9. The occurrence of patients with *LILRB3*-4SNPs among different races in multiple cohorts.**

| *LILRB3*-4SNPs | European | African American | Hispanic | Other |
|---|---|---|---|---|
| **GoCAR** (RNAseq, n = 170) [a] | | | | |
| non-Risk | 82 | 31 | 34 | 13 |
| Risk | 1 (1.2%) | 9 (22.5%) | 3 (8.1%) | 0 (0%) |
| **CTOT19** (RNAseq, n = 128) [b] | | | | |
| non-Risk | 27 | 42 | 41 | 12 |
| Risk | 0 (0%) | 5 (10.6%) | 0 (0%) | 1 (7.6%) |
| **VericiDX** (RNAseq, n=77) [c] | | | | |
| non-Risk | | 67 | | |
| Risk | | 10 (12.9%) | | |
| **Biome Biobank** (WES, n = 30,099) [d] | | | | |
| non-Risk | 9547 | 6488 | 10223 | 2950 |
| Risk | 8 (0.1%) | 608 (8.6%) | 244 (2.3%) | 31 (1%) |

* [a] genetic race; [b] genetic race; [c] self-reported race; [d] self-reported race

**Table S15. Transplant outcomes of the African American patients in the CTOT19 cohort with high expression rate of *LILRB3*-4SNPs alternative allele**

| | Expr of Ref | Expr of Alt | AEF | Race | Outcome |
|---|---|---|---|---|---|
| CTOT1912023 | 154 | 105 | 0.405 | AA | Grade 2 IF/TA, no inflammation |
| CTOT1905016 | 163 | 99 | 0.378 | AA | Poor function at 2 years (with eGFR < 30) |
| CTOT1904009 | 402 | 216 | 0.350 | AA | Normal |
| CTOT1936013 | 77 | 41 | 0.347 | AA | Poor function at 2 years (with eGFR < 30) |
| CTOT1903010 | 68 | 28 | 0.292 | AA | Grade 2 IF/TA, no inflammation |

| | Poor outcome | Normal |
|---|---|---|
| Risk | 4 | 1 |
| Non-risk | 8 | 34 |

* P = 0.01, Fisher's exact test.

**Table S16. Correlation of *LILRB3*-4SNPs with glomerulitis in AA recipients in the GoCAR cohort.**

| | Banff g <= 1 | Banff g > 1 |
|---|---|---|
| Risk | 2 | 3 |
| Non-risk | 37 | 9 |

* P = 0.044, Fisher's exact test.

**Table S17. Correlation of *LILRB3*-4SNPs with glomerulitis in AA transplant patients (ICD Z94.0) in the BioMe cohort.**

| | Banff g <= 1 | Banff g > 1 |
|---|---|---|
| Risk | 7 | 4 |
| Non-risk | 77 | 3 |

* P = 0.004, Fisher’s exact test.

**Table S18. Correlation of glomerulitis with DCGL in the GoCAR cohort.**

| | No DCGL | DCGL |
|---|---|---|
| **3-month**[a] | | |
| Banff g <= 1 | 152 | 18 |
| Banff g > 1 | 12 | 5 |
| **12-month**[b] | | |
| Banff g <= 1 | 180 | 14 |
| Banff g > 1 | 19 | 5 |
| **24-month**[c] | | |
| Banff g <= 1 | 142 | 9 |
| Banff g > 1 | 19 | 6 |

[a] P = 0.04, Fisher's exact test.
[b] P = 0.04, Fisher's exact test.
[c] P = 0.01, Fisher's exact test.

**Table S19. Comparison of sepsis (ICD code A41.9) occurrence in the AA patients with or without *LILRB3*-4SNPs or *APOL1* risk allele for kidney transplant or diseases in the BioMe cohort.**

| | Non-risk (%) | Risk (%) | Odds ratio | P value[a] |
|---|---|---|---|---|
| ***LILRB3*-4SNPs** | | | | |
| Kidney transplantation (n=130) | 13.3 (16/120) | 50 (5/10) | 6.5 | **0.006** |
| ESRD (n=465) | 13.8 (59/429) | 30.6 (11/36) | 2.76 | **0.008** |
| CKD (n=1332) | 9.3 (114/1224) | 15.7 (17/108) | 1.86 | **0.024** |
| ***APOL1* 2 Risk alleles** | | | | |
| Kidney transplantation (n=138) | 17.1 (14/82) | 11.5 (6/52) | 0.63 | 0.384 |
| ESRD (n=491) | 14.8 (53/357) | 14.2 (19/134) | 0.95 | 0.852 |
| CKD (n=1391) | 9.8 (112/1145) | 9.3 (23/246) | 0.95 | 0.835 |

[a] Logistic regression.

**Table S22. Survival analysis of *LILRB3*-4SNPs and eGFR decline (<= 15) using cox model adjusted by *APOL1* G1/G2 genotype in AA patients of the Biome cohort.**

| | 95% CI | Hazard Ratio | P value |
|---|---|---|---|
| **ESRD** (n=453) | | | |
| *LILRB3*-4SNPs | (1.05, 2.40) | 1.59 | **0.029** |
| *APOL1*-single | (0.83, 1.52) | 1.12 | 0.454 |
| *APOL1*-double | (1.09, 2.06) | 1.50 | **0.012** |
| **CKD** (n=1306) | | | |
| *LILRB3*-4SNPs | (0.81, 1.84) | 1.22 | 0.345 |
| *APOL1*-single | (0.76, 1.38) | 1.02 | 0.876 |
| *APOL1*-double | (1.88, 3.52) | 2.58 | **<0.001** |

**Table S23. Survival analysis of *LILRB3*-4SNPs alone, *APOL1*-double risk allele alone and combined (both with *LILRB3*-4SNPs risk allele and *APOL1*-double risk allele) with eGFR decline (<= 15) using cox model in AA patients of the Biome cohort.**

| | upper.95 | Hazard ratio | P value |
|---|---|---|---|
| **ESRD** (n=453) | | | |
| *LILRB3*-4SNPs alone | (1.05, 2.86) | 1.73 | **0.032** |
| *APOL1*-double alone | (1.09, 1.94) | 1.45 | **0.011** |
| Combined | (0.96, 3.99) | 1.95 | 0.066 |
| **CKD** (n=1306) | | | |
| *LILRB3*-4SNPs alone | (0.74, 2.02) | 1.23 | 0.425 |
| *APOL1*-double alone | (1.92, 3.38) | 2.55 | **<0.001** |
| Combined | (1.51, 6.26) | 3.08 | **0.002** |
